# Combined use of Oxford Nanopore and Illumina sequencing yields insights into soybean structural variation biology

**DOI:** 10.1101/2021.08.26.457816

**Authors:** Marc-André Lemay, Jonas A. Sibbesen, Davoud Torkamaneh, Jérémie Hamel, Roger C. Levesque, François Belzile

## Abstract

**Background:** Structural variant (SV) discovery based on short reads is challenging due to their complex signatures and tendency to occur in repeated regions. The increasing availability of long-read technologies has greatly facilitated SV discovery, however these technologies remain too costly to apply routinely to population-level studies. Here, we combined short-read and long-read sequencing technologies to provide a comprehensive population-scale assessment of structural variation in a panel of Canadian soybean cultivars.

**Results:** We used Oxford Nanopore sequencing data (∼12X mean coverage) for 17 samples to both benchmark SV calls made from the Illumina data and predict SVs that were subsequently genotyped in a population of 102 samples using Illumina data. Benchmarking results show that variants discovered using Oxford Nanopore can be accurately genotyped from the Illumina data. We first use the genotyped SVs for population structure analysis and show that results are comparable to those based on single-nucleotide variants. We observe that the population frequency and distribution within the genome of SVs are constrained by the location of genes. Gene Ontology and PFAM domain enrichment analyses also confirm previous reports that genes harboring high-frequency SVs are enriched for functions in defense response. Finally, we discover polymorphic transposable elements from the SVs and report evidence of the recent activity of a Stowaway MITE.

**Conclusions:** Our results demonstrate that long-read and short-read sequencing technologies can be efficiently combined to enhance SV analysis in large populations, providing a reusable framework for their study in a wider range of samples and non-model species.

## Background

Structural variants (SVs), commonly defined as genomic variations involving at least 50 nucleotides, are a key source of sequence and functional variation in eukaryotes [1–4]. Indeed, SVs such as deletions, insertions, duplications and inversions account for more variation in sequence content than single-nucleotide variants (SNVs) in several species [e.g. 5–7]. In addition to their implication in human health [8], SVs play a role in key phenotypes in crops such as soybean (*Glycine max*) [9], maize (*Zea mays*) [10, 11], tomato (*Solanum lycopersicum*) [12], wheat (*Triticum aestivum*) [13] and rapeseed (*Brassica napus*) [14]. Moreover, there is now clear evidence for the significant role played by SVs on ecological and evolutionary processes in various non-model species [15].

Despite their undeniable functional importance, genome-wide population-scale assessments of SVs have lagged behind compared to SNVs due to the lack of power of short reads for SV discovery [2]. Tools that discover SVs from short reads typically rely on one or several types of evidence, either in the form of split reads (SV breakpoint found within an individual read), discordant read pairs (unusual orientation or distance between reads of a pair), or read depth (abnormally high or low coverage at a given position) [16]. Other methods rely on local or genome-wide *de novo* assembly to discover SV breakpoints at base-pair resolution. These methods can generally detect a larger number of SVs, but they tend to struggle with repetitive SVs and on shallowly sequenced samples [17]. Unfortunately, benchmarks of tools that discover SVs from short reads consistently document sub-optimal sensitivity and precision, issues that can only be partly relieved by combining datasets obtained with different tools [18–20].

The increased availability of long-read sequencing technologies such as Oxford Nanopore and PacBio in recent years has benefited the study of SVs [21]. Indeed, their increased read length allows them to both cover the span of larger variants, such as long insertions and inversions, and to map more confidently in the low-complexity regions where SVs tend to occur [2]. Several mapping-based methods for SV discovery from long reads have already been developed [e.g. 22–25] and benchmarked [26], typically performing better than methods using short reads. These approaches have recently been applied to provide genome-wide assessments of SVs in crops such as tomato [12], rice (*Oryza sativa*) [27] and rapeseed [28].

Despite the greater power of long reads for SV discovery, their high cost and basecalling error rates make them unlikely to replace short-read technologies in the short term. In the meantime, methods that allow short-read data to use the insights gained from long reads are much needed in order to scale the study of SVs from the small cohorts sequenced with long-read technologies up to entire populations. In particular, using short-read data to genotype SVs discovered from long reads shows great promise to allow scaling up the insights gained from long reads. Although methods for genotyping SVs from short reads do exist [e.g. 29–31] and have been applied to SVs discovered from long-read sequencing data [e.g. 32], these approaches have yet to be widely adopted in plant genomics and best practices for their application in highly repetitive genomes such as that of soybean and other non-model species are still needed.

Previous studies have addressed the question of soybean structural variation using either comparative genomic hybridization [33, 34], short-read sequencing [6, 35] or pan-genome approaches [32, 36]. These studies have notably found evidence for an enrichment of SVs in genes related to defense response [33, 34, 36] and a role of SVs in determining traits such as seed coat pigmentation and iron uptake [32]. The use of a combined analysis of short and long reads could nevertheless provide new insights into soybean SV biology by allowing the study of sequence-resolved insertions efficiently and at a larger scale. Studies of transposable element (TE) polymorphisms in soybean, for example, have been limited to the identification of TE insertion boundaries [37], but long reads allow for the identification of full-length TE insertions [38].

In this study, we use an approach that combines short-read and long-read sequencing to improve prediction and genotyping of SVs in a soybean population. We first evaluate the overall performance of predicting and genotyping SVs from short reads in soybean and identify best practices for doing so. We next quantify the sensitivity and precision of genotyping Oxford Nanopore-discovered SVs using Illumina sequencing data. Finally, we combine short-read and long-read approaches to generate a comprehensive set of SVs from a panel of Canadian soybean varieties and apply this dataset to analyze population structure, relate SV location and frequency to potential impacts on gene function, and gain insights into soybean TE biology.

## Results

### Benchmarking of Illumina-discovered variants

Our first objective was to assess the performance of SV discovery and genotyping in soybean based solely on short-read sequencing data. To do this, we merged SVs discovered using four different tools (*de novo* assembly + AsmVar, Manta, smoove, and SvABA) and genotyped them in 102 samples using Paragraph. The total counts of filtered calls per discovery tool, SV type, and SV size class are summarized in Table 1. Genotype calls for 17 of the 102 samples were compared against a truth set of SVs called from Oxford Nanopore data using Sniffles and processed through a SV refinement pipeline (Additional file 1: Table S1). Comparison between Paragraph genotype calls and the ground truth was performed using the sveval R package. We only considered homozygous genotype calls for these benchmarks since we are analyzing inbred lines.

**Table 1:** Number of SVs called from the Illumina data per calling tool, SV type and size class.

| Calling program | [50bp – 100bp[ |  |  |  | [100bp – 1kb[ |  |  |  | [1kb – 10kb[ |  |  |  | ≥ 10kb<sup>a</sup> |  |  |
| --- | --- | --- | --- | --- | --- | --- | --- | --- | --- | --- | --- | --- | --- | --- | --- |
|  | DEL<sup>b</sup> | INS<sup>c</sup> | DUP<sup>d</sup> | INV<sup>e</sup> | DEL | INS | DUP | INV | DEL | INS | DUP | INV | DEL | DUP | INV |
| asmvar | 11018 | 3575 | 0 | 0 | 14877 | 2243 | 0 | 0 | 5748 | 1 | 0 | 0 | 4681 | 0 | 0 |
| manta | 9664 | 3358 | 453 | 0 | 12378 | 1815 | 3114 | 0 | 11463 | 0 | 4448 | 0 | 7325 | 5034 | 0 |
| smoove | 4168 | 0 | 22 | 45 | 6489 | 0 | 1208 | 149 | 4687 | 0 | 981 | 33 | 1794 | 975 | 47 |
| svaba | 7288 | 2284 | 673 | 21 | 6907 | 215 | 16081 | 292 | 2969 | 0 | 1548 | 190 | 512 | 458 | 223 |
| merged<sup>f</sup> | 17199 | 5023 | 656 | 61 | 22980 | 3165 | 9810 | 296 | 13007 | 1 | 4316 | 135 | 10640 | 4696 | 178 |

<sup>a</sup> Insertions ≥ 10kb are not shown because none were called <sup>b</sup> DEL: deletions, <sup>c</sup> INS: insertions <sup>d</sup> DUP: duplications <sup>e</sup> INV: inversions <sup>f</sup> merged: the dataset merged using SVmerge

Results show that the genotypes of deletions and insertions could be called confidently with as few as two (2) supporting reads, which was used as a minimum threshold for all subsequent analyses (Figure 1). At this threshold, sensitivity ranged between 50 and 65% and precision ranged from 70 to 95% for deletions, while sensitivity ranged between 30 and 40% and precision ranged from 65 to 85% for insertions (Figure 1). Precision was typically higher for intermediate-sized deletions (100-10,000 bp) than for either extremes, while sensitivity was highest for smaller ones (50-1,000 bp). Precision was higher for larger insertions than for small ones, at the expense of lower sensitivity; virtually no insertions larger than 1 kb could be called from the Illumina data (Table 1). Sensitivity increased markedly when repetitive regions were ignored, with sensitivity increasing by up to 10-20% depending on the SV type and size class, while precision remained roughly similar (Additional file 1: Figure S1). Results for inversions showed moderate precision (in the range of 40-70%) and low sensitivity (range of 10-20%), while results for duplications showed both low precision (range of 10-20%) and sensitivity (15-20%) (Additional file 1: Figure S2). Poor performance was expected for inversions and duplications given the high complexity of those types of SVs. Excluding repeat regions did little to improve the results for duplications, but it did improve sensitivity by roughly 10% for inversions (Additional file 1: Figure S3). We observed a correlation between the Oxford Nanopore sequencing depth and the genotyping precision of deletions, insertions and duplications for a given sample, with this effect being most important for duplications (Additional file 1: Figure S4). This suggests that samples that were less deeply sequenced with long reads may have failed to reveal some SVs, thus resulting in a seemingly lower precision.

**Figure 1:**
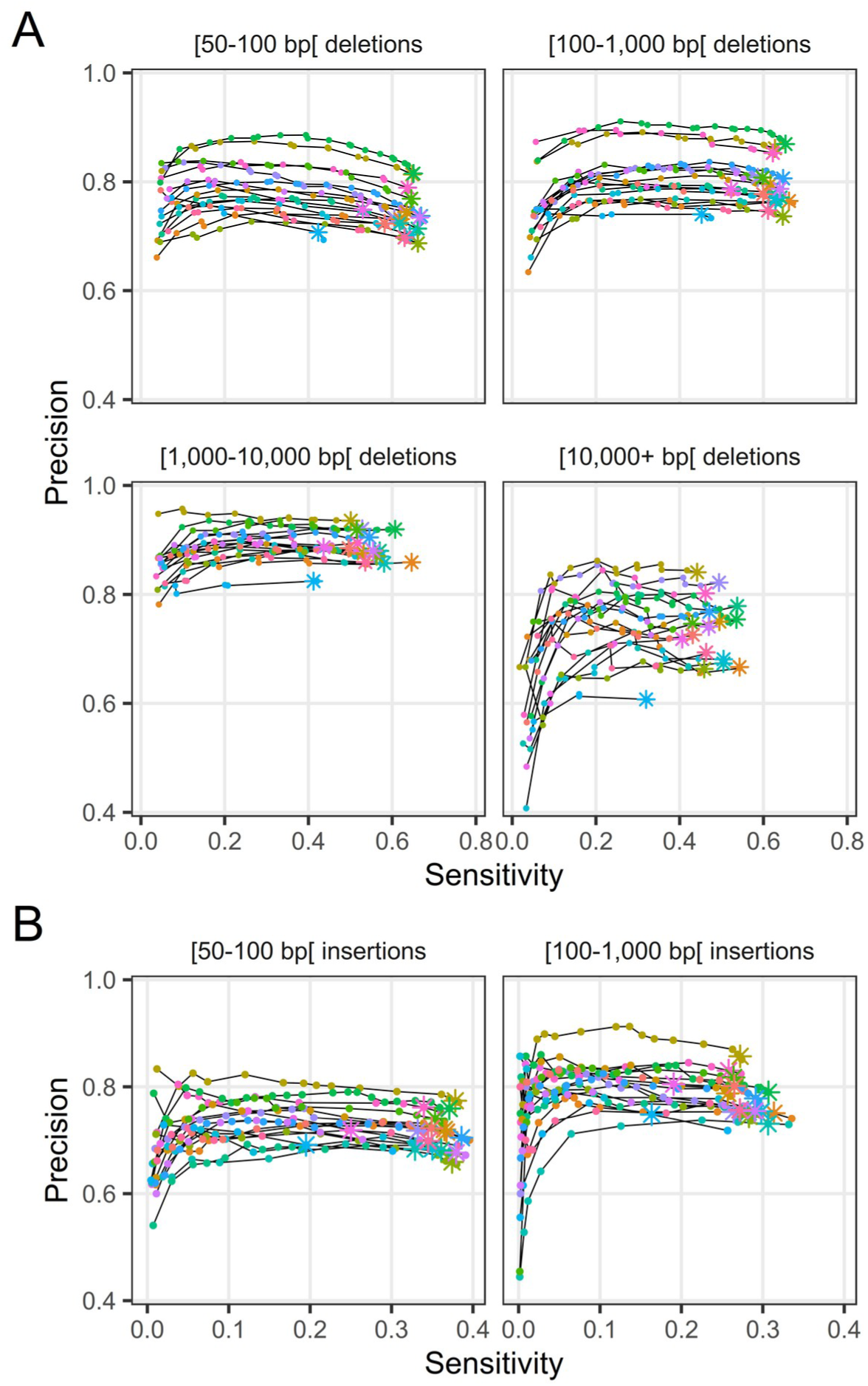
Genotyping sensitivity and precision of (A) deletions and (B) insertions discovered from the Illumina data. Each line and color represents one of 17 samples. The different plots correspond to different SV lengths. The points correspond to different filtering thresholds on the minimum number of Illumina reads required to support a genotype call. The asterisks indicate a minimum number of supporting reads of 2; points to the left of these for a given line represent increasingly stringent filtering threshold values (i.e. a greater number of reads supporting a genotype call).

Next, we assessed whether filtering SVs based on their frequency in the population resulted in a higher-quality SV set by removing putative false variants. Precision-recall curves computed for a range of homozygous ALT count (see Methods for more details) thresholds indicated that a filter based on a minimum of four alternate alleles observed across the population yielded a good compromise between sensitivity and precision for insertions and deletions (Additional file 1: Figure S5). This threshold was used to filter the set of SVs for all downstreams analyses. Filtering on the homozygous ALT count did not succeed in significantly increasing the genotyping performance of duplications and inversions (Additional file 1: Figure S6), so we decided to drop these SVs from downstream analyses. We also investigated whether a filter based on the number of distinct tools reporting a SV could be used to improve sensitivity and precision (Additional file 1: Figure S7). However, the drop in sensitivity when requiring more than one tool was generally too large to compensate for the increase in precision. Researchers valuing precision over sensitivity could however use this filter, as the gain in precision was considerable in some cases, like for large deletions.

As a consequence of their different approaches to SV discovery, and consistently with the drop in sensitivity observed when requiring multiple calling tools to consider a variant (Additional file 1: Figure S7), the various tools used showed different profiles in terms of the number of variants of different sizes and types discovered (Additional file 1: Figure S8). The performance of the different tools used for calling SVs is shown for a single representative sample in figures S9 and S10 (Additional file 1). Manta was the most important contributor of unique true positive SV calls for both deletions and insertions, followed by AsmVar. There is an obvious decrease in the false positive rate when combining evidence from several calling tools. However, individual tools still made significant contributions that justified their inclusion, with the exception of SvABA insertions which contributed few true positive SVs compared to the number of false positives (Additional file 1: Figure S10). SvABA insertions were still used for downstream analyses, but could be excluded for applications where the need for precision outweighs the need for sensitivity.

### Re-genotyping Oxford Nanopore-discovered variants

In addition to using the SVs discovered from the Oxford Nanopore data as a truth set for benchmarking SV discovery, we also assessed whether these could be accurately genotyped using Illumina data. For that purpose, we merged the calls made from the Oxford Nanopore data of all 17 samples using SVmerge. These were used as input to Paragraph and re-genotyped using Illumina data from the same 17 samples. The genotypes were compared to the SV calls made by Sniffles directly from the Oxford Nanopore data results using the sveval package as was done for the Illumina SVs.

As was the case for Illumina SVs, two (2) Illumina reads were sufficient to confidently call SVs in most samples (Figure 2). At this threshold, sensitivity ranged from 55 to 65% and precision ranged between 80 and 95% for deletions, while sensitivity ranged from 50 to 60% and precision ranged between 60 and 80% for smaller insertions (Figure 2). For deletions, sensitivity and precision were fairly consistent across size classes. For insertions, however, precision varied immensely from 20% to ∼80% for 1-10 kb insertions and from essentially 0 to 60% for insertions larger than 10 kb. Further analysis showed that there was a correlation between the precision of insertion genotyping in these size classes and the N50 of Oxford Nanopore reads of a given sample (Additional file 1: Figure S11). Therefore, it is likely that the poor precision observed for some samples is the result of limitations of the truth dataset rather than true genotyping errors. Indeed, larger insertions could not be validated in low-N50 samples because the small length of the reads prevented their discovery in those samples. Yet, those large insertions could still be genotyped using the Illumina data provided that they were discovered in other samples with higher N50. As was the case for variants discovered from the Illumina data, sensitivity was higher when we excluded repeat regions, with sensitivity reaching 80% in some cases (Additional file 1: Figure S12).

**Figure 2:**
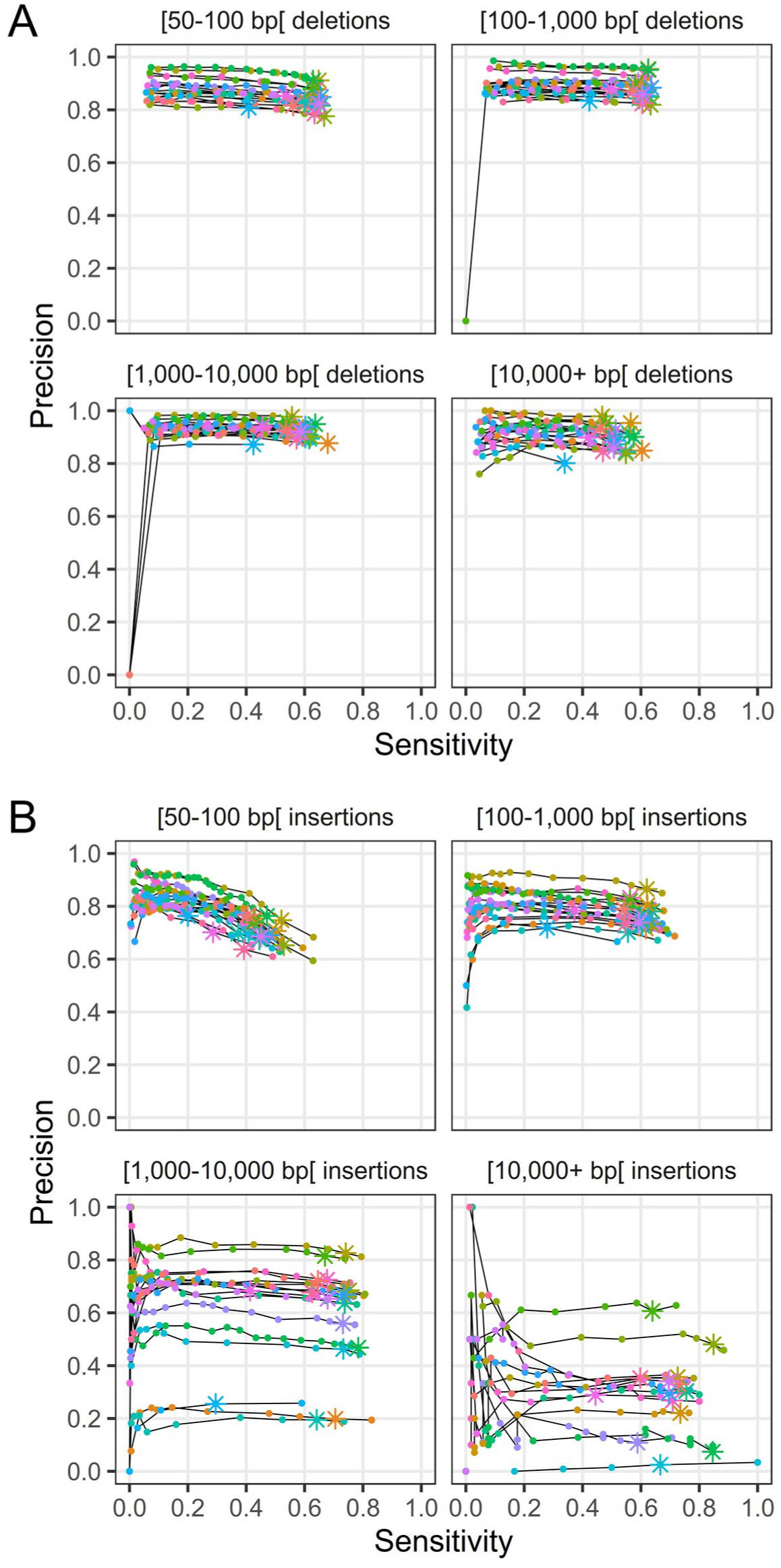
Genotyping sensitivity and precision of (A) deletions and (B) insertions discovered from the Oxford Nanopore data. Each line and color represents one of 17 samples. The different plots correspond to different SV lengths. The points correspond to different filtering thresholds on the minimum number of Illumina reads required to support a genotype call. The asterisks indicate a minimum number of supporting reads of 2; points to the left of these for a given line represent increasingly stringent filtering threshold values (i.e. a greater number of reads supporting a genotype call).

Duplications discovered by Sniffles showed low sensitivity and precision with both being in the 20-40% range (Additional file 1: Figure S13a). Inversions, however, could be accurately genotyped from the Illumina data, with a precision typically greater than 70%, but their sensitivity was low at about 10-20% (Additional file 1: Figure S13b). Concentrating on non-repeat regions moderately improved the results for duplications (Additional file 1: Figure 14a) but did so to a larger extent for inversions, with sensitivity reaching over 20% and precision being generally over 80% (Additional file 1: Figure S14b).

### Population-scale genotyping of the joint Oxford Nanopore-Illumina SV dataset

In order to produce a population-scale SV dataset that could be used for downstream analyses, we merged the SVs discovered from the Illumina and Oxford Nanopore data using SVmerge and genotyped them with Paragraph using the Illumina data of the 102 samples. Benchmarking results for deletions and insertions expectedly showed a precision that was in-between that of the previous two benchmarks (Illumina SVs and Oxford Nanopore SVs), both when considering all regions (Additional file 1: Figure S15) and non-repeat regions only (Additional file 1: Figure S16).

The dataset was further filtered using knowledge gained from the previous benchmarks. Namely, we filtered out genotype calls with fewer than two (2) supporting reads and removed SVs with fewer than four (4) alternate alleles observed among homozygous genotype calls (homozygous ALT count). Inversions and duplications were also removed for downstream analyses due to their poor performance in the benchmarks.

The distribution of deletion and insertion calls within the reference genome is illustrated in Figure 3c. There is a visible tendency for SVs to be more frequent in gene-rich euchromatic regions (Figure 3a) where predicted SNVs are also more densely distributed (Figure 3b), although this may be due only to a higher discovery power in euchromatic regions. The presence of SV hotspots on chromosomes 3, 6, 7, 16 and 18 (Figure 3c) is consistent with results previously obtained using comparative genomic hybridization by McHale et al. [34] and by a pan-genome approach [36].

**Figure 3:**
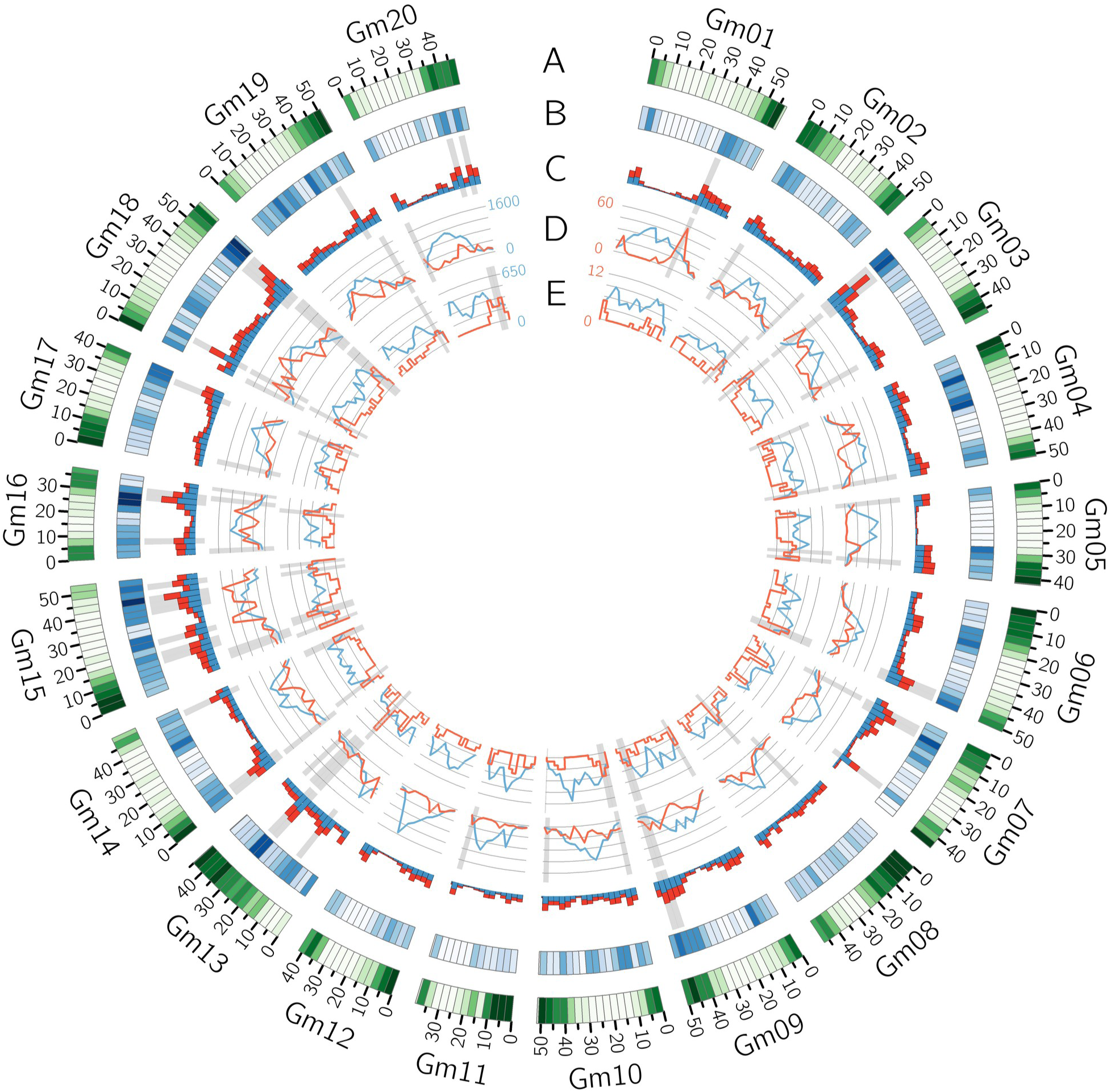
Circos plot of the distribution of various features within 3-Mb bins along the reference assembly version 4 of Williams82. (A) Gene density (B) Density of SNVs called by Platypus (C) Number of deletions (blue) and insertions (red) discovered within each bin. The bins with the 10% highest SV density (insertions and deletions considered together) are highlighted in gray. (D) Number of reference (blue) and polymorphic (red) LTR Copia and LTR Gypsy elements (summed together). (E) Number of reference (blue) and polymorphic (red) DNA transposable elements. The gray highlights in tracks D and E show the bins with the 10% highest polymorphic/reference ratios.

### Population structure

To assess the quality of our population-scale SV dataset, we verified whether population structure inferred from SVs yielded similar results to that inferred from SNVs, which are more commonly used for population structure inference. For this purpose, we assigned all individuals to one of five (5) populations using fastStructure with SNV data first, and then performed principal component analysis (PCA) on both SNV and SV data using PLINK.

The PCAs did not cluster the samples belonging to different populations into starkly distinct groups because the panel under study does not display a strong structure to begin with. Still, both the SV (Additional file 1: Figure S17a) and the SNV PCA (Additional file 1: Figure S17b) roughly grouped individuals according to their assigned population. Moreover, the PCA made from the SV genotype calls was at least as good at clustering together the samples belonging to the same population as the PCA made using SNVs was. Overall, these results constitute a proof of concept that the population-scale SV dataset is the reflection of a biological reality and not an artifact.

### Potential impact on genes

SVs can have a large impact on gene integrity or expression. Therefore, we annotated the SVs in our dataset according to the genic features they overlapped. SVs occurred disproportionately less within coding sequences than would be expected based on the proportion of the genome covered by these features, both when considering the whole genome and when restricting the analysis to non-repeat regions (Additional file 1: Table S2). A slight underrepresentation of SVs was also observed within non-coding genic sequences, although this pattern was much clearer when concentrating on non-repeat regions. Both analyses also revealed a clear pattern of overrepresentation of SVs within regions 5 kb upstream of genes. The proportion of SVs overlapping intergenic regions appeared to be less than expected when the analysis was performed on the whole genome, but this is most likely due to the fact that intergenic regions tend to be more repetitive and thus more difficult to probe. Indeed, when restricting the analysis to non-repeat regions, the proportion of SVs falling within intergenic regions was higher than their proportion within the reference genome, suggesting enrichment of SVs. We also compared the observed proportions of SVs overlapping various genic features to what would be expected by random chance using a randomization test that shuffled the positions of SVs within 100-kb bins and computed the resulting overlaps. The 100-kb bins were used to locally restrict the SVs position to take into account the repeat heterogeneity of the genome. This test confirmed the underrepresentation of SVs within coding sequences and their overrepresentation within intergenic sequences and regions 5 kb upstream of genes (Figure 4a). The pattern for non-coding genic sequences, however, diverged from other lines of evidence by suggesting slight overrepresentation of deletions. Insertions, on the other hand, appeared to be underrepresented within non-coding genic sequences, similar to the results shown in Table S2 (Additional file 1).

**Figure 4:**
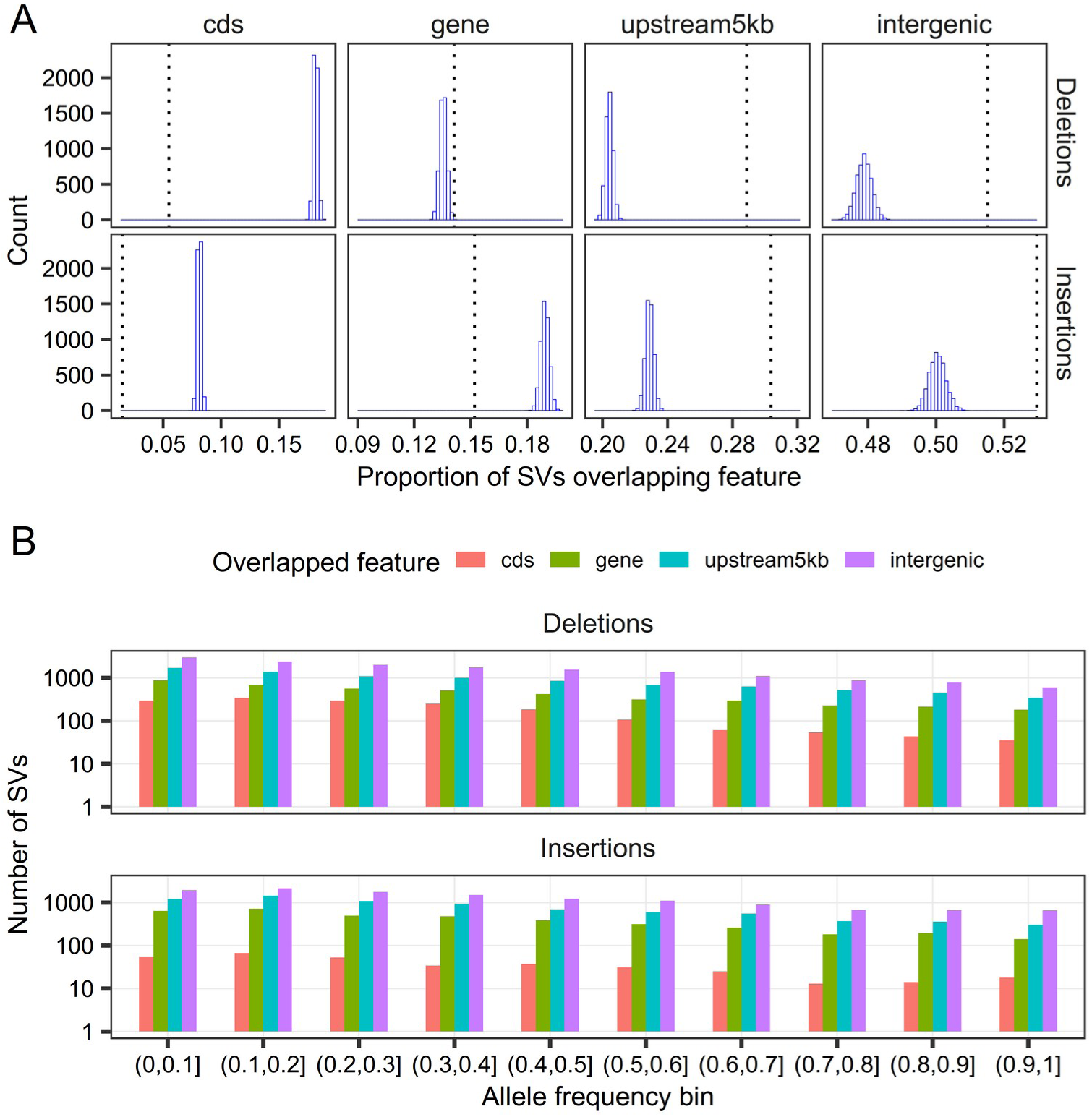
Analysis of the overlap of SVs with gene models. (A) Distributions of the proportions of deletions and insertions overlapping various genic features as generated by a randomization test (5,000 iterations). Observed proportions for each SV type and genic feature are indicated by a vertical dotted line. One-sided p-values are < 2 x 10^-4^ for all comparisons except for deletions overlapping genes, for which the p-value is 4 x 10^-4^. (B) Distribution of the allele frequencies of deletions and insertions depending on the genic features they overlap. Note the logarithmic scale on the y-axis. cds: SVs overlapping coding sequences; gene: SVs overlapping non-coding genic sequences; upstream5kb: SVs overlapping regions 5 kb upstream of genes, but not any genic sequences; intergenic: SVs that do not overlap any of the other features.

The distributions of insertion and deletion frequencies depending on the features overlapped are shown in Figure 4b. Statistical testing of the pairwise differences in mean SV frequencies depending on the genic features overlapped clearly showed that deletions overlapping coding sequences were less frequent (the frequency being lower by roughly 0.05) than those occurring elsewhere in the genome (Additional file 1: Table S3). For insertions, the only significant differences indicated a higher frequency (by roughly 0.02) in intergenic regions than in non-coding genic sequences or sequences 5 kb upstream of genes. A difference of similar magnitude was also observed between mean insertion frequency within intergenic regions and coding sequences, but the difference was marginally non-significant.

Finally, we conducted an enrichment analysis to check for over- and underrepresentation of gene ontology (GO) Biological Process terms and PFAM protein domains in genes whose coding sequence is impacted by SVs that are frequent (≥ 0.5) in the population. Genes impacted by high-frequency SVs were highly enriched for functions involved in defense response, and somewhat less so for functions involved in the regulation of various pathways (Additional file 1: Table S4; Additional file 2). Underrepresented GO Biological Process terms were almost all related to various metabolic or biosynthetic processes (Additional file 1: Table S5; Additional file 3). As was observed for GO Biological Process terms, the PFAM domain enrichment analysis showed that genes impacted by high-frequency SVs are overwhelmingly enriched in domains involved in defense response, such as NB-ARC, TIR and Leucine rich repeat domains (Additional file 1: Table S6; Additional file 4). No PFAM domains were observed to be underrepresented (Additional file 5).

### Transposable elements

Many SVs, especially larger ones, result from the mobilization of TEs [12, 39]. With this in mind, we checked whether we could gain insights into soybean TE biology from our SV dataset. To do so, we first queried the sequences of all insertions and deletions larger than 100 bp in our dataset against a database of soybean TEs. Insertions and deletions that matched a TE with high confidence were annotated with the corresponding TE type.

A total of 2,586 deletions and 2,391 insertions were annotated as TEs by this approach (Table 2; Figure 3d,e; Additional file 6). These represent 8.4% and 9.1% of all deletions and insertions, respectively, and 14.9% and 17.4% of those larger than 100 nucleotides. The proportion of polymorphic TEs of different classes found within our dataset is consistent with their prevalence in the reference genome, except for DNA TEs which represent a much smaller proportion of the polymorphic elements compared to their prevalence in the genome. The number of polymorphic elements per LTR-retrotransposon family (Figure 5a) and per DNA TE type (Figure 5b) were largely consistent with results previously reported for non-reference soybean TEs [37] except for DNA TEs of the CACTA superfamily for which we found almost no polymorphic instances.

**Figure 5:**
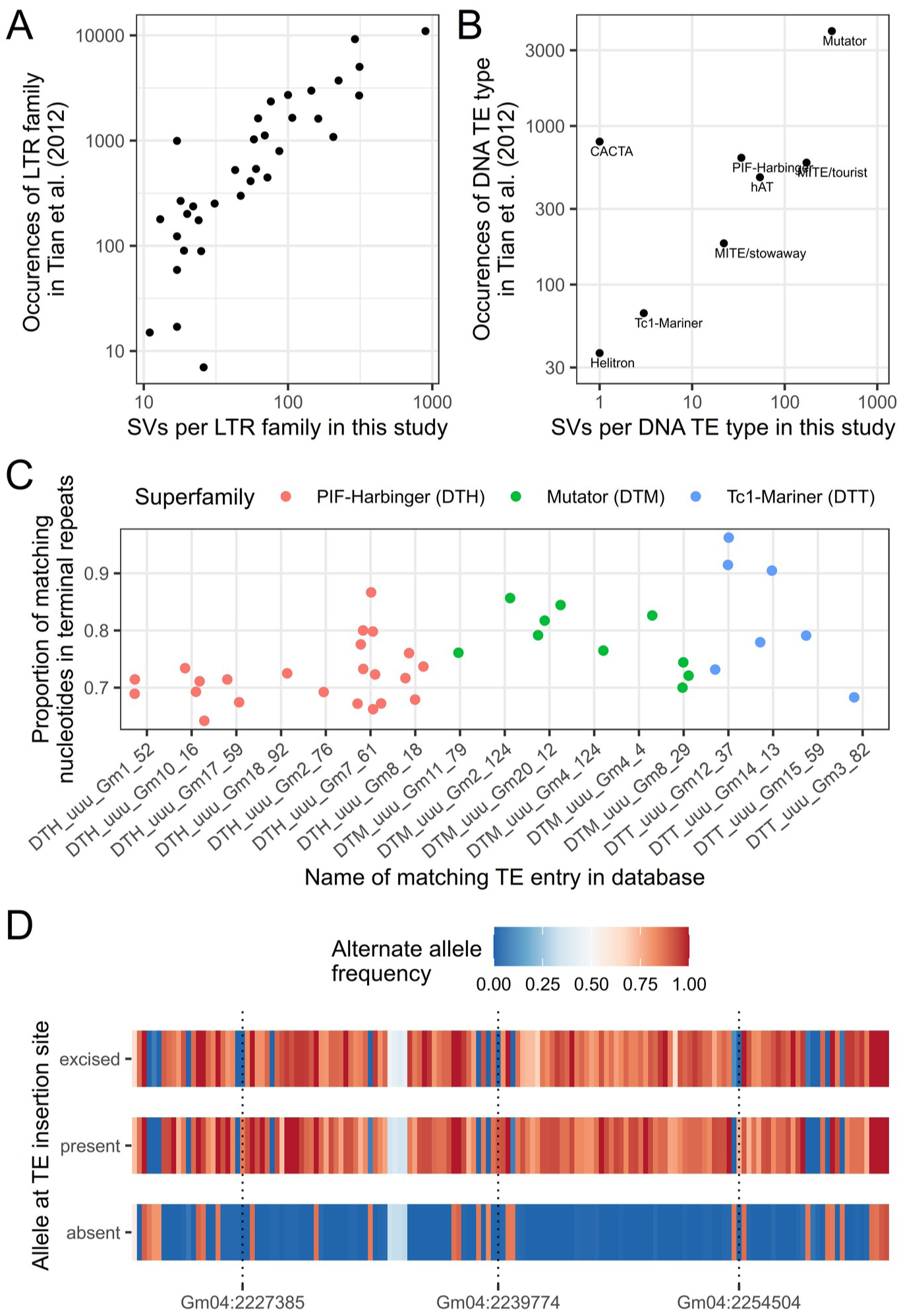
Analysis of the polymorphic TEs found in this study. Comparison of the number of polymorphic TEs per (A) LTR family and (B) DNA TE type found in Tian et al. [37] and in this study. Differences in y- and x-scales are partly explained by the fact that counts for Tian et al. are summed over occurrences in all samples whereas our data counts each SV only once. Note that all scales are logarithmic. (C) Proportion of matching nucleotides between the two terminal repeats for TE sequences corresponding to 40 different SVs grouped by DNA TE superfamily and by the identifier of the TE sequence they matched in the SoyTEdb database. (D) Alternate allele frequencies of 156 SNVs located in a ∼39-kb linkage disequilibrium block between positions Gm04:2,220,398 and Gm04:2,259,326. Frequencies were computed for three different groups of samples depending on their genotype at the TE insertion site (Gm04:2,257,090). absent: absence of the TE insertion, which corresponds to the reference allele (71 samples); present: presence of the 480-bp Stowaway MITE (9 samples); excised: presence of a 6-bp insertion at the insertion site, putatively left by excision of the TE insertion (14 samples). The locations of three SNVs whose frequency in the “present” and “excised” groups diverge are shown with dotted vertical lines.

**Table 2:** Number and span of polymorphic and reference transposable elements of different types.

| TE type | REF<sup>a</sup>(%) |  | DEL<sup>b</sup>(%) |  | INS<sup>c</sup>(%) |  |
| --- | --- | --- | --- | --- | --- | --- |
|  | *N*<sup>d</sup> | Mb<sup>e</sup> | *N* | kb<sup>f</sup> | *N* | kb |
| Copia LTR retrotransposons | 91241 (35.1) | 170 (43.0) | 1154 (44.6) | 5594 (43.8) | 1303 (54.5) | 6692 (63.1) |
| Gypsy LTR retrotransposons | 71390 (27.5) | 139 (35.2) | 949 (36.7) | 5745 (45) | 718 (30) | 2949 (27.8) |
| Non-LTR retrotransposons | 8078 (3.1) | 10 (2.5) | 144 (5.6) | 449 (3.5) | 99 (4.1) | 307 (2.9) |
| DNA TE | 89300 (34.3) | 76 (19.2) | 339 (13.1) | 989 (7.7) | 271 (11.3) | 654 (6.2) |

<sup>a</sup> REF: transposable elements ≥ 100 bp in the reference genome <sup>b</sup> DEL: deletions relative to the reference that are annotated as TEs <sup>c</sup> INS: insertions relative to the reference that are annotated as TEs <sup>d</sup> *N*: Number of reference elements, deletions or insertions matching given TE type <sup>e</sup> Mb: Total length of reference elements of a given type, in Mb <sup>f</sup> kb: Total length of polymorphic elements matching given TE type, in kb

We identified terminal inverted repeat (TIR) and target site duplication (TSD) sequences from local assemblies of the TE sequences for 40 different polymorphic SVs collectively representing 17 entries in the SoyTEdb database. The polymorphic TEs for which we could identify TIR and TSD sequences were essentially miniature inverted-repeat transposable elements (MITEs) ranging in size from 198 to 681 bp (longer sequences were too challenging to assemble properly). From these data, we computed the proportion of matching nucleotides between the two inverted repeats and averaged the values over the all samples bearing the TE insertion for a given SV (Figure 5c). A high proportion of matching nucleotides can indicate the potential for active transposition because intact transposons should have identical or nearly identical TIRs. While the proportion of matching nucleotides was under 0.8 in most cases, three polymorphic TEs matching the Tc1-Mariner superfamily and annotated as Stowaway MITEs presented a proportion of matching nucleotides > 0.9 (Additional file 7).

We generated multiple alignments of the local assemblies at all sites where at least one sample had recognizable TIR and TSD sequences. A visual analysis of these multiple alignments revealed that for all but one SV, the sequences that did not bear the insertion presented a single occurrence of the TSD sequence. This observation is consistent with a scenario where the TE never inserted into the sequence, instead of having excised from it. The one exception to this observation is that of a 480-bp insertion of a Stowaway MITE at position 2,257,090 of chromosome Gm04. In this case, a visual analysis of the multiple alignment revealed that three different alleles are segregating in the population at the insertion site: (1) the reference allele (no insertion at the target position), (2) a 480-bp insertion that corresponds to the TE insertion, and (3) a 6-bp insertion of nucleotides TACGAG (Additional file 1: Figure S18; Additional file 8). Interestingly, this insertion is by far the one for which the percent similarity between the two TIR sequences was highest among the ones studied, at 96.3%. We hypothesized that the 6-bp insertion resulted from the excision of the TE, with the TA nucleotides being remnants of the classical Tc1-Mariner TSD and the other nucleotides having been added during DNA repair following excision. If this is the case, then the haplotypes surrounding the insertion site should be very similar between the individuals with the TE insertion and those with the 6-bp insertion. Using a combination of SV calls made by Paragraph and indel calls made by Platypus, we assigned 71 individuals as homozygous for the reference allele, 9 individuals as homozygous for the TE insertion allele and 14 individuals as homozygous for the 6-bp insertion allele. We computed the alternate allele frequencies within each of these three groups for 156 SNVs located in a 39-kb linkage disequilibrium block surrounding the insertion site (Figure 5d). The results clearly show high genetic similarity between individuals bearing the TE insertion and those bearing the 6-bp insertion, consistent with the latter being derived from excision of the TE insertion. In fact, only three (3) SNVs showed contrasting allele frequencies (difference in allele frequencies > 0.5) between these two groups (Figure 5d), whereas 129 alleles were contrasted between the reference allele haplotype and the TE insertion allele haplotype. This suggests that the excision of the TE is a relatively recent event and that this TE may still be active in soybean. Interestingly, one of the polymorphic *Copia* insertions found in our dataset matches an insertion in the Glyma.20G090000 gene (also known as the PhyA2 gene corresponding to the E4 maturity locus) known to impact time to maturity in soybean [40]. In our dataset, this TE insertion had a frequency of 0.207, with 20 samples genotyped as homozygous for the alternative allele and a single one genotyped as heterozygous.

## Discussion

The rapid development of long-read sequencing platforms such as PacBio and Oxford Nanopore in recent years has greatly enhanced the potential for studying structural variation. Although studies using long reads to survey structural variation in crops have started to emerge [e.g. 12, 27, 28], they did not explicitly address the question of using short reads to scale up SV analysis from the small cohorts sequenced using long reads to larger populations, as has been done in humans [e.g. 41, 42]. This question is of interest because long-read sequencing remains too expensive at the moment to apply at large scale and because large amounts of already-existing short-read sequencing data could be leveraged in that way. Scaling up the study of SVs is a necessary prerequisite to getting a clear understanding of genome evolution and function, and applying this knowledge to real-world problems [1, 15]. In this study, we demonstrate that a relatively small cohort of 17 samples sequenced to ∼12X coverage with Oxford Nanopore can be combined with Illumina data to drive the study of SVs in a population of 102 Canadian soybean lines and gain insights into SV biology.

The SVs discovered from short-read sequencing data are typically limited to variants located in non-repeated regions and relatively small insertions (< 200 bp). These results have been shown repeatedly by benchmarking studies [18, 20] and reflect an inherent limitation of short reads to span large repeats and effectively assemble into long insertions. Here, we still used Illumina reads for SV discovery to survey the whole population and thus detect less frequent variants that may not have been found within the 17 samples sequenced with Oxford Nanopore data. However, despite following recommended practices for SV analysis such as combining different SV calling tools and integrating the results with a dedicated SV genotyper, estimated sensitivity for insertions remained low at ∼40% for those in the range 50-100 bp and ∼30% for those in the range 100-1,000 bp. The improved sensitivity obtained when focusing on non-repeated regions (up to ∼60% for insertions in the range 50-100 bp) shows that a large part of the problem indeed comes from repeated regions. However, entirely removing these regions from analyses is an unsatisfactory solution as polymorphisms in these regions may still be relevant to a particular study question.

To compensate for limitations in SV discovery from short reads, we assessed whether Illumina reads could be used to genotype SVs discovered from Oxford Nanopore data on a smaller cohort of 17 samples. The greatest added value of this approach arguably comes from the possibility to accurately genotype large (> 1 kb) insertions with > 70% sensitivity. This is an encouraging result because it shows that such insertions can be successfully genotyped using Illumina data even though they could not be discovered from this same data. This is because long reads provide the full contiguous sequence of insertions, which the Illumina reads can then map to. Combined with a novel pipeline for refining the breakpoints and sequence content of SVs discovered from Oxford Nanopore sequencing data prior to genotyping, this approach should enable the study of SVs in large populations for which short-read data is already available. The main limitation to using this approach actually comes from the long-read data itself. In this study, some of the samples appeared to have low genotyping precision for larger insertions, but this was most likely due to these insertions not being discovered in samples with lower read N50 and thus appearing as false positive genotype calls. Similarly, the sequencing depth of the Oxford Nanopore data used here was not sufficient to provide a solid reference dataset for benchmarking duplications. Indeed, one limitation of our study is that Oxford Nanopore reads alone do not provide a perfect ground truth for benchmarking, especially for SVs under 100 bp [2], but this was the best truth set we had access to in the absence of a gold standard SV dataset for soybean.

Follow-up analysis on our population-scale SV dataset confirmed that this dataset reproduced previously described population structure patterns, an validation approach commonly used in other population-scale SV studies [e.g. 43, 44]. We indeed found that a PCA using SVs summarized the population structure just as well as a PCA using SNVs, which indicates that the SV genotype calls on the 102-sample population are accurate. Perhaps more importantly, the SV dataset produced here met our expectations regarding the genome-wide distribution of SVs and their location relative to predicted gene models. The location of SV hotspots found here is consistent with previously reported results [34, 36]. Moreover, GO term and PFAM domain enrichment analyses confirmed previous observations that SV-enriched genes were involved in plant defense response [33, 34, 36]. Several lines of evidence in our results also suggest a strong functional constraint on the location of SVs in the soybean genome. Notably, SVs were strongly depleted within coding sequences compared to what would be randomly expected, and insertions were depleted within non-coding genic sequences. There was also a clear tendency for enrichment of SVs in regions upstream of genes, but whether this is simply due to lower functional constraints or a role of SVs in regulating gene expression remains to be investigated. Functional constraints on the frequency of SVs could also be observed from our data, as deletions impacting coding sequences were less frequent than those occurring elsewhere in the genome and insertions were enriched within intergenic regions, which are arguably less functionally important. Based on these results, we suggest that many of the deletions located within coding sequences may have a deleterious impact and could therefore become targets for breeding.

The large insertions and higher power of SV discovery within repetitive regions that was afforded by the Oxford Nanopore sequencing data gave us an opportunity to study soybean TE biology more deeply than previous reports. The numbers of TEs associated with various superfamilies was largely consistent with results previously reported by Tian et al. [37], except for DNA TEs of the CACTA superfamily which were a lot less common in our data. We observed the same pattern of general concordance with previously reported results except for CACTA elements when comparing our data to that of Istanto [45]. The reason why we found almost no polymorphic CACTA elements compared to these studies is unclear, but we hypothesize that it may be due to our more stringent requirements for TE annotation. Indeed, we required the length of the queried SVs to be close to that of their matching counterpart in the database. Many of the SVs in our dataset indeed matched CACTA elements following the BLASTN query, but almost all of them failed to pass the filter. Our annotation results are probably conservative for other types of TEs as well because the database we used is likely incomplete, as it is based on the analysis of a single reference genome.

Our data also allowed us to generate original findings related to DNA TEs in soybean, which have received relatively little attention from past studies. We report results that suggest that most DNA TE insertion polymorphisms in soybean result from past insertion of TEs rather than from excision of existing TEs. The relatively low proportion of polymorphic DNA TEs compared to their prevalence in the genome also suggests that these elements are overall fairly inactive in soybean. However, we did document one case in which recent excision of a Stowaway MITE from its insertion site appears to have occurred, such that three alleles (the reference allele without the insertion, the TE insertion, and the allele resulting from the excision of the TE) are present within the population. This element represents a prime candidate to study the potential activity of DNA TE transposons in soybean.

## Conclusions

In conclusion, our study shows that Oxford Nanopore and Illumina sequencing data can be efficiently combined to study structural variation in soybean. In particular, large insertions that cannot be discovered from short-read data alone could be genotyped using short-read data and thus allow the insights gained from long-read sequencing to scale up to a larger population. This approach, combined with a novel pipeline for refining the SVs discovered using Oxford Nanopore data, should extend easily to other species and allow the wealth of already-existing Illumina data to be leveraged for SV analysis. In addition to confirming previous results regarding the chromosomal distribution of SVs in soybean and their association with genes involved in defense response, we also report novel insights into functional constraints to the occurence of SVs and into soybean TE biology. Moreover, the SV catalog described here is freely available and can be used as a resource for SV genotyping by the soybean research community. These results as well as the framework developed to optimize the study of structural variation at population scale should help to better integrate these variants in genomic studies of crops and other non-model species.

## Methods

### Illumina sequencing and read processing

Sample selection and acquisition of Illumina sequencing data has been described in previous work [6]. Briefly, 102 Canadian soybean cultivars and breeding lines were selected to encompass the full range of genetic variation found among Canadian short-season germplasm and sequenced on the Illumina HiSeq 2500 platform. Paired-end reads ranging in size from 100 to 125 nucleotides were obtained depending on the sample. This sequencing data is available on the NCBI Sequence Read Archive (SRA) through BioProject accession number PRJNA356132 [46].

All reads were adapter- and quality-trimmed using bbduk from the BBtools suite v. 38.25 [47]. We aligned reads using bwa mem v. 0.7.17-r1188 [48] with default parameters. Paired-end alignment mode was used except for reads that were left unpaired following adapter and quality trimming, which were aligned in single-end mode. We used a reference genome consisting of assembly version 4 of the Williams82 reference cultivar [49] concatenated with reference mitochondrion and chloroplast sequences retrieved from SoyBase [50]. Reads aligned using paired-end and single-end mode were then merged, sorted and indexed using samtools v. 1.8 [51] and read groups were added using bamaddrg [52]. The sorted and indexed BAM files were used as input for all downstream analyses requiring mapped reads.

### Structural variation discovery from short reads

We called SVs on all 102 samples using four different tools: AsmVar [53], Manta [54], SvABA [55], and LUMPY-based [56] smoove [57]. We selected this combination of tools based on the complementarity of their SV detection approaches, widespread use within the community, and performance reported in published benchmarks [20].

AsmVar calls SVs by comparing *de novo* genome assemblies to a reference genome. Prior to assembly, we merged reads that were still paired after trimming using FLASH v. 1.2.11 [58]. The rationale behind this was that the short size of the inserts in our sequencing data allowed several of the read pairs to be merged into longer sequences. Reads were grouped into three libraries (single-end reads from bbduk, single-end reads merged by FLASH, and paired-end reads left unmerged by FLASH) and assembled with SOAPdenovo2 v. 2.04 [59] using the sparse_pregraph and contig commands, and a k-mer size of 49. Contigs were not further assembled into scaffolds because we aimed to only call SVs whose sequence was entirely resolved. The resulting contigs were aligned to the reference genome using LAST v. 1047 [60] by first calling the lastal command with options -D1000 -Q0 -e20 -j4 and then the last-split command with options -m 0.01 -s30. Variants were called on the LAST alignments using ASV_VariantDetector from the AsmVar tool suite (version of 2015-04-16) with default parameters. The pipeline was run on each sample independently and results were subsequently concatenated to obtain a single AsmVar VCF file. Variants with a FILTER tag other than “.” were filtered out from the resulting call set.

We ran manta v. 1.6.0 with default parameters in 10 batches of 10 or 11 randomly grouped samples because it did not scale well to the whole population. We used the candidate SVs (and not the genotype calls themselves) identified by each run for further processing and filtered them by removing unresolved breakends (SVTYPE=BND). The filtered variants were then converted from symbolic alleles (i.e. DEL, DUP, INS) to sequence-explicit ALT alleles using bayesTyperTools convertAllele v. 1.5 [31] and combined into a single VCF file using bcftools merge (version 1.10.2-105) [51].

We ran SvABA v. 1.1.3 separately on all samples using the command svaba run with options -- germline -I -L 6. SvABA produces two different variant sets: one for indels, which are already coded as sequence-explicit, and another for SVs which are coded as paired breakends. We therefore classified SVs into defined types (DEL, DUP, INV) based on breakpoint orientation and converted them to sequence-specific ALT alleles using an in-house R script. The resulting sequence-explicit variants were merged using bcftools merge.

We ran smoove v. 0.2.4 on all samples using a series of commands. First, smoove call was run separately on each sample using default parameters. The variants identified were then merged into a single VCF file using smoove merge, smoove genotype with options -x -d, and smoove paste. Symbolic alleles (<del>, <DUP> and <INV> alleles) were converted to explicit sequence representation using bayesTyperTools convertAllele.

A series of common filters were applied to the SV output of all four tools before using them for downstream analyses. Specifically, we removed variants spanning less than 50 bp or more than 500 kb, those located on unanchored scaffolds or organellar genomes, or any variant that was not classified as either a deletion, insertion, duplication or inversion. We also converted multiallelic variants into biallelic records and standardized the representation of all alleles using bcftools norm.

### Oxford Nanopore sequencing

We selected 17 samples for Oxford Nanopore sequencing among those sequenced by Illumina. Sixteen (16) of them were randomly selected among a subset of 56 lines belonging to a core set of Canadian soybean germplasm, while the remaining sample (CAD1052/OAC Embro) had been selected and sequenced before the others based on its higher Illumina sequencing depth. Although sample selection did not explicitly maximize the number of potential SVs assessed, we did verify that the resulting set covered the range of variation found in Canadian soybean germplasm based on an existing phylogenetic tree [6].

Our sample preparation and sequencing protocols evolved throughout the project as we gained experience with Oxford Nanopore sequencing. Therefore, we outline our latest methods here, but more details regarding the procedures used for each sample can be found in Table S7 (Additional file 1). Accessions selected for sequencing were germinated in Jiffy peat pellets (Jiffy Group, Zwijndrecht, Netherlands) on the benchtop. Young trifoliate leaves were collected between two and three weeks after germination, flash frozen in liquid nitrogen upon harvest and stored at -80 °C until DNA extraction. Single trifoliate leaves weighing between 20 and 60 mg were used for each extraction. Liquid nitrogen-frozen leaves were pulverized on a Qiagen TissueLyser instrument (Qiagen, Hilden, Germany) with metal beads for four cycles of 30 s each at 30 Hz. The resulting powder was immediately transferred to a CTAB buffer (2% CTAB, 0.1 M Tris-HCl pH 8, 0.02 M EDTA pH 8, 1.4 M NaCl, 1% (m/v) PVP) and incubated at 60°C in a water bath for 45 min. The lysate recovered after centrifugation at 3500 rcf for 10 minutes was then subjected to an RNase A treatment for another 45 min at 60°C, followed by the addition of an equal volume of 24:1 chloroform:isoamyl alcohol to the sample and stirring to an emulsion. Following centrifugation at 3500 rcf for 15 minutes, the supernatant was recovered and mixed with a 0.7 volume of cold isopropanol. This mix was stored at -80°C for 20 minutes and centrifuged at 3500 rcf for 30 min, after which the liquid was removed. Tubes were rinsed twice with cold 70% ethanol, with a centrifugation step after each addition of ethanol. After the last rinsing, tubes were left to dry for 3 minutes after which pellets were resuspended in 100 µl elution buffer (Tris-HCl 0.01 M and EDTA 0.001 M, pH 8) at 37°C for an hour, and then stored at 4°C until use.

Samples were size-selected using the Short Read Eliminator kit of Circulomics (Circulomics, Baltimore, MD, USA) following the manufacturer’s instructions. The size-selected DNA resuspended in the SRE kit’s EB buffer was then purified using SparQ magnetic beads and resuspended in ddH_2_O. Typically, between 500 ng and 1 µg of this DNA was used for Oxford Nanopore library preparation using the SQK-LSK109 genomic DNA ligation kit (Oxford Nanopore Technologies, Oxford, UK). The library was prepared according to the manufacturer’s instructions except for the following details: 1) DNA fragmentation was not performed prior to library preparation, 2) 80% ethanol was used instead of 70% ethanol, 3) the bead elution time following DNA repair and end-prep was increased from 2 min to 10 min, 4) the bead elution time following adapter ligation and clean-up was increased from 10 to 15 minutes and carried out in a water bath set to 37°C. Typically, between 150 ng and 400 ng of the prepared library quantified using a Qubit fluorometer (Thermo Fisher Scientific, Waltham, MA, USA) were used as input to a FLO-MIN106D flowcell (R9 chemistry) and run on a MinION for 48 to 72 hours using default voltage settings. While most accessions were sequenced on a single flow cell, three accessions for which the initial yield was low (< 9 Gb) were sequenced a second time (using DNA from a different plant) to provide sufficient data for downstream analyses. More details regarding the Oxford Nanopore sequencing of the samples can be found in Table S8 (Additional file 1).

### Structural variation discovery from Oxford Nanopore data

Raw FAST5 sequencing files were basecalled on a GPU using Oxford Nanopore Technologies’ guppy basecaller v. 4.0.11 with parameters --flowcell FLO-MIN106 --kit SQK-LSK109. Basecalled FASTQ files obtained from a single flow cell were concatenated into a single file which was used for downstream analyses. Adapters were trimmed using Porechop v. 0.2.4 [61] with the option --discard_middle. Adapter-trimmed reads were aligned using NGMLR v. 0.2.7 [24] with the option -x ont. The resulting alignments were sorted and indexed using samtools.

At this stage, we merged the BAM files of samples that were sequenced on two different flowcells and called SVs using Sniffles v. 1.0.11 [24]. We ran Sniffles with parameters -- min_support 3 (minimum number of reads supporting a variant = 3, default = 10), -- min_seq_size 1000 (minimum read segment length for consideration = 1000, default = 2000) and --min_homo_af 0.7 (minimum alternate allele frequency to be considered homozygous = 0.7, default 0.8). We chose relaxed parameters compared to the defaults because our samples are inbred cultivars and heterozygosity should therefore be nearly non-existent.

We applied a series of filters to the SVs in order to remove any spurious calls that could affect downstream analyses. Any variants called on organellar genomes or unanchored scaffolds were filtered out, along with any variants smaller than 50 nucleotides or larger than 500 kb. We only retained deletions, insertions, inversions and duplications for further analyses, discarding unresolved breakpoints (SVTYPE=BND) as well as other complex types such as DEL/INV, DUP/INS, INVDUP and INV/INVDUP. We removed variants called as heterozygous since heterozygous genotype calls are very likely to be spurious in these inbred lines. In order to avoid calling artificial variants in ambiguous regions of the genome (stretches of “N” due to imperfectly assembled regions of the reference genome), we also removed deletions that overlapped any “N” in the reference as well as any insertion located less than 20 nucleotides away from any “N” in the reference.

The location of SVs as well as the insertion sequences reported by Sniffles are necessarily imperfect as they are based on error-prone Oxford Nanopore reads (on average 8-10% error rate based on the percent identity of our alignments). We therefore assembled a pipeline to refine the breakpoint location and the sequence content of the deletions and insertions found by Sniffles. Duplications and inversions were not considered for SV refinement because the inherent complexity of these variants made it difficult to accurately assemble them from our data. We briefly describe the pipeline here, but more details can be found in Additional file 1 (Supplemental Methods, Table S9 and Figures S19 to S21). Our breakpoint refinement pipeline starts by locally assembling all reads that were mapped by NGMLR to positions ± 200 bp from the location of the SV using wtdbg2 v. 2.5 [62]. The same reads are then aligned to the assembled sequence using minimap2 v. 2.17-r974 [63] to polish the assembly sequence using the consensus module of wtdbg2. The resulting polished assembly is subsequently aligned to the local region of the reference genome using AGE (commit 6fa60999, github.com/abyzovlab/AGE) [64]. The coordinates of the SV and insertion sequence content are then optionally updated from the information provided by the AGE alignment. When the alignment did not suggest suitable replacement coordinates or insertion content for a given SV, we simply used its representation as initially defined by Sniffles for downstream analyses instead. Following breakpoint refinement, the representation of the alleles was standardized using bcftools norm.

### Structural variant genotyping and benchmarking

We genotyped SVs on all 102 Illumina samples using Paragraph v. 2.4a [29] in three different batches. The first batch used only variants discovered from the Illumina data as input and was used to assess the performance of SV discovery from Illumina data alone. The second batch used only variants discovered from Oxford Nanopore data and was similarly used to assess the performance of genotyping those variants with Illumina data. The third and last genotyping batch used a merged dataset comprising both variants discovered using Illumina and Oxford Nanopore data, and was used for the population-scale analyses on population structure, location of variants relative to gene models, and polymorphic TEs. Despite the superior performance of long-read data for SV discovery, we decided to also include variants discovered from the Illumina data in the final SV set as they encompassed all samples.

For genotyping SVs discovered from Illumina data, the VCF files of all discovery tools (AsmVar, Manta, SvABA, smoove) were merged together using SVmerge (commit 6a18fa3d2, github.com/nhansen/SVanalyzer) [65] with parameters -maxdist 15 -reldist 0.2 -relsizediff 0.1 - relshift 0.1. Parameters were chosen in order to merge slightly differing representations of alleles that were putatively identical from a biological point of view while preserving true allele diversity at a given position.

SVs discovered from Oxford Nanopore data were also merged across samples using SVmerge with the same parameters as described above. However, for Oxford Nanopore variants, we modified SVmerge’s default behavior which selects an allele randomly from a given SV cluster. Instead, we forced the random selection to be made among the alleles that had been refined by the SV refinement pipeline, if any, to favor those alleles whose representation was hopefully closer to biological reality.

For the last batch combining Illumina and Oxford Nanopore variants, the two datasets described above were merged using SVmerge. The default behaviour of SVmerge was again overridden by systematically sampling among the alleles found by Illumina whenever a SV cluster contained alleles found by both Illumina and Oxford Nanopore. Despite the greater power of Oxford Nanopore data in discovering SVs, our reasoning was that if a variant was discovered by both sequencing technologies, then the Illumina data was likely more precise given its higher basecalling accuracy.

The methods used for genotyping were identical for all three batches. We prepared the VCF files for input to Paragraph by removing variants located less than 1 kb away from chromosome ends and padding the allele representations as required by Paragraph. We genotyped the 102 Illumina samples aligned by bwa mem following the recommendations outlined by Paragraph for population-scale genotyping, i.e. the variants were genotyped independently for each sample with multigrmpy, setting the -M option to 20 times the average sequencing depth for the sample.

We compared the genotyping results of the three batches against the Oxford Nanopore SV set in order to assess genotyping sensitivity and precision. For this analysis, the set of variants called from the Oxford Nanopore data by Sniffles and subsequently refined was considered to be the ground truth. Structural variation calls made from Oxford Nanopore data may also be erroneous, especially for smaller variants [2], so this approach of treating Oxford Nanopore dataset as the ground truth is necessarily imperfect but nevertheless provides a good comparison basis for our purposes.

We compared the SV genotype calls to the ground truth set using the R package sveval v. 2.0.0 [30]. For each of the 17 samples for which Oxford Nanopore data was available, we compared the genotype calls made by Paragraph to the SVs identified in the Oxford Nanopore data for that sample. SVs genotyped as homozygous for the alternate allele by Paragraph and present in the Nanopore set were considered true positives, while SVs genotyped as homozygous for the alternate allele by Paragraph but absent from the Nanopore set were considered false positives. Note that, for benchmarking purposes, we essentially ignored heterozygous genotype calls made by Paragraph since the truth set only contained homozygous calls as expected for inbred lines. Sensitivity was defined as the ratio of the number of true positive calls to the total number of SVs in the truth set, and precision as the ratio of the number of true positive calls to the sum of true and false positive calls. We computed sample-wise precision-recall curves for various SV size classes and SV types by using a range of read count thresholds (number of reads required to support a genotype call) to filter the Paragraph genotype calls. We required sveval to explicitly compare insertion sequences by setting ins.seq.comp = TRUE, but we otherwise used default settings. We extended sveval’s functionality by also assessing duplications under the same overlap conditions as the package already provides for deletions and inversions. Benchmarks were performed both on the complete set of SVs and on a subset of SVs located in non-repeat regions. A SV was defined as belonging to a repetitive region if it had a 20% or higher overlap to regions in the repeat annotation for the Williams82 assembly version 4 retrieved from Phytozome [66].

For the SVs discovered by Illumina, we computed additional precision-recall curves by filtering the SVs in the dataset genotyped by Paragraph based on two different metrics of SV quality: (1) the number of times the alternate allele is observed in homozygous genotype calls across the whole population (referred to hereafter as the homozygous ALT count) and (2) the number of calling tools (out of a maximum of four) that originally reported the SV. The more stringent homozygous ALT count was used instead of alternate allele frequency as a measure of the frequency of the SV in the population since true SVs are expected to be homozygous for the alternate allele in these inbred lines. Note that both of these quality measures (homozygous ALT count and the number of tools supporting an SV) effectively filter SV records and not individual genotype calls. The objective of these analyses was to see whether filtering on SV frequency or calling-tool support for variants could result in a higher quality dataset.

### Population structure

We used the set of merged Illumina and Oxford Nanopore SVs genotyped by Paragraph to evaluate whether SV calls could replicate population structure analyses made from SNV calls. We applied methods similar to Torkamaneh et al. [6] in order to compute population structure for the 102-sample population. We called SNVs using Platypus v. 0.8.1.1 [67] with parameters -- minMapQual=20 --minBaseQual=20 --maxVariants=10 --filterReadsWithUnmappedMates=0 -- filterReadsWithDistantMates=0 --filterReadPairsWithSmallInserts=0. We filtered Platypus calls to keep only biallelic SNVs located on any of the 20 reference chromosomes. We only retained SNVs with a minor allele frequency ≥ 0.05, proportion of missing sites ≤ 0.4, and heterozygosity rate ≤ 0.1. The resulting 1.27 M SNVs were converted to PLINK BED format [68] and used as input to fastStructure v. 1.0 [69] using k = 5 as determined by Torkamaneh et al. [6]. A PCA was computed on those SNVs using PLINK v1.90b5.3 with default parameters. A PCA was also computed on the population-scale dataset of Illumina/Oxford Nanopore SVs genotyped with Paragraph. For this analysis, we filtered SV genotype calls by setting those with less than two supporting reads to missing. We also removed duplications, inversions, as well as records with a homozygous ALT count < 4 or a proportion of missing sites ≥ 0.4.

### Potential impact on genes

We annotated deletions and insertions based on their overlap with various gene features. We retrieved the positions of the gene models for Williams82 assembly 4 from Phytozome [66] and determined for each SV whether it overlapped any of the following genic features: coding sequences, non-coding genic sequences, and regions 5 kb upstream of genes. These categories were mutually exclusive, such that an SV overlapping both coding and non-coding sequences was only labeled as “coding sequences”. Similarly, an SV was only labeled as “5 kb upstream” if it did not overlap any genic sequences. The SVs that overlapped none of the features described above were labeled as “intergenic”.

We first used these annotations to assess whether SVs were over- or underrepresented within particular genic features by comparing the observed proportions of deletions and insertions overlapping each feature to what would be expected by chance. We used three different measures of random expectation of the proportion of SVs overlapping genic features. The first measure was a naive comparison to the proportion of the genome corresponding to each genic feature. This comparison is however biased because repetitive regions (which are largely non-genic) are less effectively queried for SVs than non-repetitive genic regions. Therefore, we also replicated the analysis by excluding repeated regions, which provided a second measure of random expectation. Finally, we performed a randomization test by estimating the distribution over the proportions of SVs that would be expected to overlap each genic feature by random chance. This was done by shuffling the start positions of SVs within the 100-kb genome-tiling bins in which they are located 5,000 times and annotating them with the genic features overlapped. We used 100-kb bins tiled along the whole genome instead of shuffling the positions genome-wide to take into consideration the heterogeneity of the genome while allowing SVs to be repositioned in a gene-agnostic manner.

We also used the genic feature annotations to study differences in mean alternate allele frequencies of SVs depending on the features they overlapped. We averaged the frequencies of insertions and deletions overlapping each of the four genic features and computed the difference between the mean SV frequencies for each of the six possible pairwise combinations of features. SVs with a frequency of 1 in the population were excluded from this analysis because they might be due to errors in the reference assembly. Statistical significance was assessed using a randomization test by shuffling the genic feature annotations 10,000 times to get a distribution of mean SV frequency differences between feature groups under a random scenario. We computed one-sided *p*-values by comparing the observed values to the random distributions thus generated, using a significance threshold of α = 0.05 / 6 = 0.0083 to compensate for multiple testing.

Finally, we carried out enrichment analyses of GO [70] Biological Process terms and PFAM domains [71] to assess whether high-frequency gene-impacting SVs were associated with particular biological functions. We identified insertions and deletions with an alternate allele frequency ≥ 0.5 and < 1 among those overlapping coding sequences and found 546 genes overlapped by such SVs. These genes constituted our gene set of interest for the enrichment analyses. We used the GOstats Bioconductor package v. 2.56.0 [72] along with GO and PFAM annotations for Williams82 assembly version 4 retrieved from Soybase on April 20 2021 to test this gene set for over- and underrepresentation of particular GO Biological Process terms or PFAM protein domains. We only tested GO terms and PFAM domains that were represented by at least 20 and 10 genes, respectively. For the GO terms, we used the conditional test as implemented in GOstats and the GO.db annotation package v. 3.12.1 [73]. We applied a Bonferroni correction to the p-values of both the GO and PFAM enrichment tests by multiplying the p-values by the number of terms/domains tested.

### Transposable elements

We annotated TEs in the SVs discovered using the SoyTEdb database [74] downloaded from SoyBase [50]. We queried the deleted or inserted sequences of all deletions and insertions ≥ 100 bp against SoyTEdb using blastn v. 2.11.0+ [75] with default parameters. Any queried sequence that aligned to a TE in the database with at least 80% of the query length and 80% of the length of the TE sequence was considered a match and annotated accordingly with the classification of the best-matching TE. All alignments that matched these criteria had an extremely small E-value (< 10^-80^) and therefore no additional filtering on this was needed.

The annotated SVs were then used to determine both the proportion of polymorphic TEs belonging to each category and the physical location of polymorphic TEs in the genome. We also computed the proportions of TEs ≥ 100 bp in each category within the reference repeat annotation from Phytozome and compared those to the estimated proportions in the SV dataset. The estimated number of polymorphic TEs within various LTR-retrotransposon families and DNA TE types were also compared to the number of non-reference TEs found by Tian et al. [37] to check whether our results were consistent with previous reports.

Soybean DNA TEs have received little attention compared to retrotransposons, which are more prevalent and polymorphic in this species [37, e.g. 76]. DNA TEs that have TIR typically transpose using a “cut and paste” mechanism. This mechanism generates a TSD upon insertion into the genome, and leaves this TSD as well as possible additional nucleotides upon excision due to DNA repair [77]. In order to study the dynamics of polymorphic DNA TEs within our population, we devised a pipeline based on local assembly and multiple sequence alignment of the DNA TE insertions. Briefly, the pipeline locally assembles Oxford Nanopore reads surrounding the sites of polymorphic DNA TEs for all samples using wtdbg2 and aligns these assemblies to each other using MAFFT v. 7.475 [78] before identifying TIR and TSD sequences with Generic Repeat Finder v. 1.0 [79]. For more details on the pipeline, see Supplemental Methods (Additional file 1). Our goal with this pipeline was to determine whether the insertion/deletion polymorphisms at various sites were due to novel TE insertion, TE excision, or a combination of both phenomena. We applied this pipeline to SVs that were annotated as TIR DNA TEs and whose matching sequence in the SoyTEdb database was matched by at least three SVs. We limited ourselves to TE sequences that were matched by at least three SV events under the assumption that TEs present in multiple copies were more likely to have been recently active. For insertions that had both TIR and TSD sequences unambiguously identified, we computed the proportion of matching nucleotides in the alignment of the two terminal repeats and averaged the values across all local assemblies bearing the insertion in order to get a single value for that SV.

### Software used

Unless otherwise stated, all statistical analyses and data manipulation were conducted in R version 3.5.0 or 4.0.3 [80] and Bioconductor version 3.08 or 3.12 [81]. Analyses made use of Bioconductor packages Biostrings v. 2.58.0 [82], GenomicRanges v. 1.42.0 [83], Rsamtools v. 2.6.0 [84], rtracklayer v. 1.50.0 [85] and VariantAnnotation v. 1.36.0 [86]. All scripts used for the analyses described in this paper are available on GitHub [87].

## Additional files

Additional file 1: Supplemental methods, supplemental tables S1 to S9 and supplemental figures S1 to S21 (PDF 12 MB)

Additional file 2: Statistics of the conditional hypergeometric test for the overrepresentation of GO Biological Process terms using the GOstats R package. P-values are Bonferroni-corrected p-values. (CSV 141 KB)

Additional file 3: Statistics of the conditional hypergeometric test for the underrepresentation of GO Biological Process terms using the GOstats R package. P-values are Bonferroni-corrected p-values. (CSV 265 KB)

Additional file 4: Statistics of the hypergeometric test for the overrepresentation of PFAM domains using the GOstats R package. P-values are Bonferroni-corrected p-values. (CSV 18 KB)

Additional file 5: Statistics of the hypergeometric test for the underrepresentation of PFAM domains using the GOstats R package. P-values are Bonferroni-corrected p-values. (CSV 95 KB)

Additional file 6: SVs identified as polymorphic transposable elements among the dataset of combined Illumina/Oxford Nanopore variants genotyped with Paragraph. Positions of the SVs and metadata about their best blastn match in the SoyTEdb database are described. (CSV 598 KB)

Additional file 7: Proportion of matching nucleotides in TIR of SVs for which intact TSD sequences and matching TIR were identified with GenericRepeatFinder (CSV 2.4 KB).

Additional file 8: Multiple alignment of the Williams82 assembly version 4 reference sequence and local *de novo* assemblies of 7 samples at the site of a 480-bp Stowaway MITE insertion (Gm04:2,257,090). Samples OAC Petrel and Roland bear the 480-bp insertion, while Alta bears the 6-bp TACGAG insertion; other samples match the reference sequence. Asterisks mark the locations of the TSD sequences in samples OAC Petrel and Roland. (TXT 6.2 KB)

## Abbreviations

GO: gene ontology
MITE: miniature inverted-repeat transposable element
PCA: principal component analysis
SRA: sequence read archive
SV: structural variant
SNV: single- nucleotide variant
TE: transposable element
TIR: terminal inverted repeat
TSD: target site duplication

## Declarations

### Ethics approval and consent to participate

Not applicable.

### Consent for publication

Not applicable.

### Availability of data and materials

The Illumina sequencing data used in this study is available on the SRA repository through BioProject accession number PRJNA356132 [46]. The Oxford Nanopore sequencing data generated during the current study is available in the SRA repository through BioProject accession number PRJNA751911 [88].

The SoyTEdb database is available on SoyBase [89].

Gene Ontology annotations for Williams82 assembly version 4 [90] as well as chloroplast and mitochondrion genome sequences [91] are also available on SoyBase.

The non-reference transposable elements found by Tian et al. (2012) can be downloaded from the supplementary data to their paper [37].

The reference genome sequence and annotation of soybean cultivar Williams82, assembly version 4, are available on Phytozome [92].

VCF files generated during this study and results of the permutation test for the analysis of the proportion of SVs overlapping various genic features are available on the figshare repository [93].

The code for all the analyses described in the paper [87] and the breakpoint refinement pipeline [94] are available on GitHub. Versions of the code at the time of submission are archived on the figshare repository for the breakpoint refinement pipeline [95] and the analysis code [96].

### Authors’ contributions

Conception and design of the study: MAL, JAS, DT, FB. Illumina sequencing: DT. Oxford Nanopore sequencing: MAL, JH, RCL. Data analysis: MAL. Data interpretation: MAL, JAS, FB. Manuscript drafting: MAL, JAS, FB. All authors have revised the manuscript and approved its submission.

## Acknowledgements

The authors thank Brian Boyle for assistance with Oxford Nanopore sequencing, Maxime de Ronne for providing the CTAB protocol, Anders Krogh for hosting MAL at the University of Copenhagen, Claire Mérot for providing comments on the manuscript, Malcolm Morrision for providing some of the plant materials, as well as Dave Deandre Istanto and Matthew Hudson for providing access to unpublished data on soybean polymorphic transposable elements. None of the funding bodies were involved in study design, data acquisition, data analysis, interpretation of the results, or manuscript writing.

