## Additional file 1 for "Combined use of Oxford Nanopore and Illumina sequencing yields insights into soybean structural variation biology"

#### Supplemental methods

**1 Description and testing of the SV refinement pipeline** **4**

| | | |
|---|---|---|
| Figure S11 | Relationship between Oxford Nanopore read N50 and genotyping precision of Oxford Nanopore insertions | 33 |
| Figure S12 | Sensitivity and precision of Oxford Nanopore deletion and insertion genotyping in non-repeat regions | 34 |
| Figure S13 | Sensitivity and precision of Oxford Nanopore duplication and inversion genotyping | 35 |
| Figure S14 | Sensitivity and precision of Oxford Nanopore duplication and inversion genotyping in non-repeat regions | 36 |
| Figure S15 | Sensitivity and precision of joint Illumina/Oxford Nanopore deletion and insertion genotyping | 37 |
| Figure S16 | Sensitivity and precision of joint Illumina/Oxford Nanopore deletion and insertion genotyping in non-repeat regions | 38 |
| Figure S17 | Population structure analyses of the whole Canadian soybean panel | 39 |
| Figure S18 | IGV screenshot of Oxford Nanopore and Illumina read alignments of three samples at the location of a polymorphic Stowaway TE insertion | 40 |
| Figure S19 | Genotyping sensitivity of raw and refined Oxford Nanopore SVs for four samples using BayesTyper | 41 |
| Figure S20 | Genotyping sensitivity of raw and refined Oxford Nanopore SVs for four samples using vg | 42 |
| Figure S21 | Genotyping sensitivity of raw and refined Oxford Nanopore SVs for four samples using Paragraph | 43 |

### 1 Description and testing of the SV refinement pipeline

One of the main objectives of this project is to use Illumina data to genotype SVs that were originally discovered using Oxford Nanopore sequencing data. This would enable the use of Illumina data to carry out population-scale genotyping of variants that were initially discovered in a smaller set of samples sequenced with long-read technologies.

Oxford Nanopore data, however, has a high error rate. This makes it challenging to accurately describe breakpoints and sequence content (for insertions) and may negatively impact our ability to genotype the variants discovered with this technology using short reads.

In order to address this issue, we assembled a pipeline that can be used to refine the SVs discovered by Sniffles [1]. By refinement, we mean that the position of the breakpoints (start and end position for deletions, and single location for insertions) and the sequence content (for insertions) can be updated to improve the quality of the variants. The current version of the pipeline ignores SVs other than deletions or insertions due to difficulties in assembling and aligning them properly.

The pipeline is based purely on Oxford Nanopore data (i.e. it does not need Illumina data e.g. for polishing or error correction) and therefore could potentially be used for SV refinement with any Oxford Nanopore or even PacBio dataset.

#### 1.1 Details of the steps in the pipeline

The pipeline performs a local assembly of the Oxford Nanopore reads aligned to the region of interest and then uses AGE [2] to align this assembly to the appropriate region of the reference genome. This alignment is then used to update the breakpoints and inserted sequence of the variant if certain quality thresholds are met.

This alignment pipeline is largely inspired by the guidelines suggested by the authors of AGE in their newer LongAGE paper [3] and the authors of wtdbg2 [4] and consists of the following steps:

1. For every insertion or deletion, Oxford Nanopore reads overlapping the region of interest are extracted from the NGMLR [1] BAM file. The region of interest is a window spanning ± 200 nucleotides from the SV breakpoints (start and end of the breakpoint for deletions, and single location for insertions).
2. The reads extracted in this way are assembled using wtdbg2. We use preset values for Oxford Nanopore sequencing (`-x ont`) except for the following parameters: `-S 1` (no subsampling of reads), `-e 2` (minimum read coverage of 2 for an edge in the graph), `-L 1000` (minimum read size of 1 kb), `-g 500000` (expected genome size of 500 kb).

3. A consensus sequence for the assembly is produced using the `wtpoa-cns` command from wtdbg2 with default parameters.

4. This assembly is polished using Oxford Nanopore data. For this, the same reads that were used for assembly are aligned to the consensus sequence using minimap2 [5] with options `-ax map-ont -r2k`. These aligned reads are used to update the consensus sequence with `wtpoa-cns`.

5. Following assembly and polishing, the reference sequence is extracted ± 500 nucleotides from the SV breakpoints for the assembled contig to be aligned against it.

6. The largest of the assembled contigs (there is usually only one) is then aligned to the reference sequence using AGE with parameters `-allpos`, `-go=-1` (gap opening penalty of -1 to account for the high indel error rate of Oxford Nanopore data), `-indel` (the expected excised region is an indel) and `-both` (try alignment of both the sequence and its reverse complement, and report the best alignment).

Once the alignment is completed, the results are parsed and used to extract the location of the regions excised by AGE from both the assembled contig and the reference genome. In the case of a deletion, we expect a fragment of the reference genome to be excised, whereas in the case of a insertion, we expect a fragment of the assembled contig to be excised. From this information, the location of the excised fragment is transformed to reference-based coordinates and the inserted sequence is extracted from the assembled contig (using the reverse complement if applicable). The following metrics are then computed in order to compare the (refined) deletion/insertion suggested by the AGE alignment to the original (raw) one defined by Sniffles:

- For deletions, the reciprocal overlap between the raw deletion and its refined counterpart. The reciprocal overlap is the minimum of the two following values: proportion of deletion 1 covered by deletion 2, and proportion of deletion 2 covered by deletion 1. This overlap is computed using functions from the GenomicRanges R package [6].

- For insertions, the offset of the two insertions, i.e. the (absolute) physical distance in base pairs between the positions of the raw and refined insertion.

- For insertions, the Levenshtein (edit) distance between the two inserted sequences, as computed by the R function `adist`. From this distance, a *distance ratio* is computed, which corresponds to the Levensthein distance between the two insertions divided by the length of the longest insertion. This provides a relative measure of the difference in sequence content between the two insertions.

- In addition to the metrics related to the excised fragments themselves, the percentage of nucleotides that share identity and the percentage of nucleotides affected by gaps in the aligned flanking regions are also extracted from the output of AGE.

Note that SVs longer than 50 kb are excluded from the pipeline because of the high memory and computing time requirements of aligning to those variants using AGE. Although a newer version of AGE called LongAGE [3] requiring less memory has been recently released, we found it unsuitable for our needs due to cryptic error messages and slower speed.

#### 1.2 Parameters determining whether SVs are updated or not

Our objective is to update the position of the breakpoints and the sequence content such that the SVs updated in this way are closer to the actual sequence, and thus both easier to genotype from Illumina data and more representative of reality. However, we must also avoid updating the variant in a way that is inaccurate (as would occur if, for example, AGE identified a deletion or insertion different from the target one). Therefore, we established conditions to decide whether or not an SV should be updated. Based on analyses carried out during the development of the pipeline, we have settled on the following thresholds to decide whether a variant should be updated:

- Reciprocal overlap $\geq 0.5$ (for deletions)
- Offset $\leq 50$ (for insertions)
- Edit distance ratio $\leq 0.5$ (for insertions)
- Percent identity of the alignment $\geq 85$
- Percentage of gaps in the assembly $\leq 15$

#### 1.3 Testing the effect of refined SVs on genotyping performance

The main objective of the SV refinement pipeline described above is to improve the genotyping of SVs with Illumina data. In order to verify whether the pipeline indeed improved genotyping, we compared the genotyping precision and sensitivity of raw and refined SVs using three different genotypers: BayesTyper v. 1.5 [7], vg v. 1.23.0 [8] and Paragraph v. 2.4a [9].

##### 1.3.1 Application of the pipeline

For these testing purposes, we applied the SV refinement pipeline described above to the first four samples for which we obtained Oxford Nanopore data. SVs discovered using Sniffles were filtered as described in the main text (notably, only SVs genotyped as homozygous were kept) and then used as input to the pipeline. Insertions and deletions were refined if they met the required thresholds, which were as described above except for the minimum reciprocal overlap which was set to 0.7 and the maximum distance ratio which was set to 0.45.

Table S9 shows the number of variants that could be refined or not for each sample. For all four samples, a majority of SVs subjected to the pipeline met the conditions for refinement.

There can be three main reasons why a variant is not refined:

- The proposed refined variant did not meet one or several quality thresholds when compared to the raw variant.
- The size of the variant was > 50 kb.
- No assembly or alignment was produced by the pipeline.

In either of these cases, we simply keep the original representation of the SV as called by Sniffles.

###### 1.3.2 Input SV datasets

In order to assess the effect of refining SVs on genotpying performance, we generated an input dataset of raw variants and another input dataset of refined variants.

In order to generate input datasets for genotyping, the SV sets obtained from the four samples were merged using SVmerge [10] with parameters `-maxdist 15 -reldist 0.1 -relsizediff 0.1 -relshift 0.1 -seqspecific`. For refined variants, the default random selection behavior of SVmerge when combining several variants together was modified to systematically favor variants that had been updated by the pipeline.

Following this, the VCF files were coordinate-sorted, normalized (using `bcftools norm`, [11]) and duplicate records were removed.

###### 1.3.3 Genotyping methods

The raw and refined SV datasets were both used as input to three genotyping programs, BayesTyper, vg and Paragraph.

**BayesTyper** We followed the methods suggested by the authors of BayesTyper for genotyping (`https://github.com/bioinformatics-centre/BayesTyper`).

Briefly, k-mers were counted from the adapter-trimmed fastq files using KMC [12] and a Bloom filter was applied using the `bayesTyperTools makeBloom` utility.

In parallel, `BayesTyperTools combine` had to be run to make the VCF input files suitable for input to BayesTyper.

Genotyping was then carried out using `bayesTyper cluster` and `bayesTyper genotype` sequentially with default parameters except for the use of the mitochondrial and chloroplastic genome sequences as decoy sequences.

Precision-recall curves for BayesTyper were generated by varying the genotype quality (GQ) threshold necessary to make a call.

**vg** Variation graphs were created from the VCF files using `vg construct` and then converted to PackedGraph format using `vg convert -p`. The graphs were then indexed prior to mapping using `vg index -L -x` to generate the xg index and `vg index -g` to generate the GCSA index. GCSA indexing was carried out on a version of the graph that had been pruned with `vg prune`.

Reads were mapped separately for each sample using `vg map`. These mappings were then used to call the SVs explicitly defined in the input VCF files using `vg pack -Q5` and `vg call -v`.

Precision-recall curves for vg were generated by varying the genotype quality (GQ) threshold necessary to make a call.

**Paragraph** VCF input files were prepared for input to Paragraph using similar methods to those described in the main text, i.e. variants located less than 500 nucleotides away from chromosome ends were removed and variants were padded with reference nucleotides where required. Samples were genotyped separately using the `multigrmpy` utility of Paragraph.

Precision-recall curves for Paragraph were generated by varying the minimum number of reads (DP) threshold necessary to make a call.

##### 1.3.4 Benchmarks: raw versus refined variants

Benchmarks were carried out using the sveval R package [8, `https://github.com/jmonlong/sveval`] following the methods described in the main text.

**Bayestyper** Figure S19 shows the benchmarking results obtained for BayesTyper. Overall, the refined deletions resulted in a higher sensitivity but similar precision. The benefits of SV refinement were obvious for larger deletions, whereas results were sometimes worse for deletions in the range of 100-1000 nucleotides. The improvement brought by refinement is more obvious for insertions than it is for deletions, most likely because the refinement pipeline also improved the sequence content of the insertions. The poor overall sensitivity of the results obtained with BayesTyper when genotyping Oxford Nanopore SVs is probably due to the small tolerance of this program to errors in SV breakpoint location or sequence content, as has been previously documented [8]. Nevertheless, the improvement in sensitivity obtained by using refined SVs suggests that the SV refinement pipeline indeed improves the quality of the variants.

**vg** Figure S20 shows the benchmarking results obtained for vg. Refining the deletions does seem to have improved the results, although only very slightly. For insertions, however, the improvement resulting from refinement is obvious, both in terms of sensitivity and precision. Again, this suggests that the breakpoint refinement pipeline improved the accuracy of the sequence content of insertions, such that they could be more accurately genotyped by vg.

**Paragraph** Figure S21 shows the benchmarking results obtained for Paragraph. Using refined variants results in a slight increase in sensitivity for Paragraph, both for deletions and smaller insertions ($<$ 1,000 bp). The apparent drop in precision for insertions $\geq$ 1,000 bp is likely due to the fact that the Oxford Nanopore data represented a poor ground truth dataset for these samples in this size range, as explained in the main text. It is therefore expected that an increase in sensitivity from refining the variants will result in an apparent loss of precision in this size range.

#### 1.4 Conclusion on the SV refinement pipeline

Overall, the results show that the breakpoint refinement pipeline results in higher genotyping sensitivity than using the raw variants as called by Sniffles. While the results are not immediately obvious when using Paragraph, the improvements obtained using BayesTyper and vg suggest that the quality of the SVs is indeed higher following breakpoint refinement. This is probably due to the high tolerance of Paragraph to errors in SV sequence (as documented in [9]) compared to the two other tools. However, even if the improvement in genotyping performance using Paragraph is not obvious, downstream analyses should still benefit from more accurate SV breakpoints and sequence.

#### 1.5 Software availability

The SV refinement pipeline described here is available from `https://github.com/malemay/breakpoint_refinement`.

### 2 Detailed methods on the DNA transposable element analysis

#### 2.1 Description of the analysis pipeline

To study potential DNA transposable element (TE) activity in more detail, we extracted the SVs corresponding to all DNA TEs in the SoyTEdb database [13] that were matched by at least 3 SVs. This represented 51 TEs in the database and 237 individual SVs.

For each of these SVs, we extracted the Oxford Nanopore reads at the location of the SV and assembled them individually for each sample following methods similar to those used for the breakpoint refinement pipeline. Briefly, reads overlapping positions ± 200 bp from the SV were used for assembly with wtdbg2 and polishing with wtpoa_cns after re-aligning the reads to the assembly with minimap2.

The assemblies were then used for multiple alignment across samples in order to identify TE boundaries as well as target site duplication (TSD) and terminal inverted repeat (TIR) sequences. We used the `PairwiseAlignment` function from the Bioconductor Biostrings package [14] to individually align all assemblies to a section of the reference genome ± 500 bp from the putative TE insertion site. This pairwise alignment was only done in order to extract the part of the assemblies that was of interest.

We then used the aligned sequences obtained from the pairwise alignment as input for multiple alignment. Multiple alignment was done with the MAFFT [15] suite of tools using the command `ginsi --reorder` which performs multiple global alignment.

We processed multiple alignments so as to only keep high-quality alignments that showed obvious TE insertion polymorphism. First, we manually checked all alignments and only kept those that showed an insertion/deletion polymorphism of the expected length. Second, for a given multiple alignment, several of the sequences often showed very poor similarity to the reference genome, possibly because of issues with their assembly. We removed such sequences by filtering out those that had less than 75% percent identity with the reference in the first and last 500 bp of the alignment (the nucleotides flanking the insertion/deletion site). Typically, only short TEs (< 500 bp, essentially MITEs) remained following these filtering steps because the longer ones were too challenging to assemble and align properly.

A total of 72 SV events representing 25 individual entries in the TE database remained following filtering. The sequences that remained following these filtering steps typically differed only in the presence or absence of the TE sequence. Sequences that contained the TE sequence could be analyzed further to identify TSD and TIR sequences from the TE insertion sequence. 70 out of the 72 SVs had at least one sequence bearing the insertion after filtering. In total, 301 sequences bearing the insertion could be queried for TSD and TIR sequences. 31 of these 301 were extracted from insertions that are present in the reference, whereas the 270 remaining insertion sequences were extracted from the local assemblies of the Oxford Nanopore reads.

We analyzed the insertion sequences using the GenericRepeatFinder software [16] with the following command:

```
grf-main -i input_fasta -o .  -c 1 -p 30 -s 10 --seed_mismatch 8 --min_tr 12
```

The parameters `--min_space` and `--max_space` were adjusted to be -60 and +20 relative to the expected SV size, respectively. The minimum (`--min_tsd`) and maximum (`--max_tsd`) TSD length were set to 2 for the DTT superfamily and 3 for the DTH superfamily, while these values were 5 and 10, respectively, for the DTM superfamily. We used relatively relaxed parameters for seed matching because the presence of TSD and TIR sequences was expected and we wanted to maximize the probability of finding them.

We parsed the output of GenericRepeatFinder to get the location and sequences of the TSD and TIR. In many cases, several potential TSD/TIR sequences could be identified from a given input sequence. For DTT TEs, only entries with a TSD matching the expected TA sequence were kept. For DTH TEs, only entries with a TSD matching either TAA or TTA were kept. Despite those filters, several TSD/TIR candidates often remained for a given sample. Therefore, the candidates were ranked following an automated procedure according to their proximity to the expected insertion location and similarity to expected insertion size. For TEs in the DTM family, the highest priority was given to longer TSDs. For every sample, we considered the TSD/TIR entry with the highest priority as the only match for that sample.

We annotated the multiple alignments with the putative TSD/TIR sequences and visually checked for consistency between these sequences and the location of the insertion. Our analysis relied entirely on the automatically identified TSD/TIR sequences; samples for which the best matching TSD/TIR sequence was not consistent were discarded and were not analyzed further. In total, we identified high-quality TSD/TIR sequences for 42 individual SVs. For each of the TIR sequences identified, we computed the percentage of matching nucleotides between the two inverted repeats and averaged them over all insertion sequences for a given SV.

Some of the TIR sequences were identified directly from the reference sequence and may thus have been of higher quality than the error-prone assemblies generated from the Oxford Nanopore data. Therefore, we only show the results obtained from the de novo-assembled sequences in figure 5c of the main text. This results in 40 individual SVs collectively representing 17 sequences in the SoyTEdb database.

#### 2.2 Detailed analysis of a 480-bp Stowaway MITE insertion

A closer look at the multiple alignments revealed that in almost all cases, the sequences without the insertion presented a single occurence of the TSD sequence. This observation is consistent with a scenario where the TE never inserted into the sequence, instead of having excised from it. In a single case however, that of a 480-bp insertion of a Tc1-Mariner (DTT) element (actually a Stowaway MITE) at position 2,257,090 of chromosome Gm04, visual analysis of the multiple alignment revealed that three different alleles are segregating at the insertion site:

- The reference allele (no insertion at the target position)
- A 480-bp insertion that corresponds to the TE insertion
- A 6-bp insertion of nucleotides TACGAG

We speculate that this last allele results from the excision of the TE insertion. Indeed, the TA part of the insertion corresponds to the expected TSD sequence for a Tc1-Mariner element. The remaining CGAG nucleotides would likely have been inserted during the repair of the DNA break following excision of the TE. Interestingly, this insertion is by far the one for which the similarity between the two TIR sequences was highest among the ones studied, at 96.3%.

To investigate this question further, we analysed the SNV patterns surrounding the insertion site. If the TACGAG indeed resulted from the excision of the TE, then the haplotypes corresponding to the TE insertion and the 6-bp insertion should be more similar to each other than to the reference sequence, as the excision would have occurred within the TE insertion genetic background.

The SNVs with FILTER = PASS among those discovered by Platypus [17] were extracted from the VCF file of the 102 samples sequenced by Illumina to perform this analysis. The analysis of linkage disequilibrium (LD) using PLINK [18] in a 200-kb window surrounding the insertion location at Gm04:2,257,090 revealed a high-LD block spanning positions 2,220,398 to 2,259,326 that included the location of the insertion. The SNVs in that 39-kb window were therefore extracted from the Platypus VCF file and used for downstream analyses.

By combining the information obtained from Platypus and SV genotyping with Paragraph, we were able to classify samples into three groups depending on their status at the TE insertion locus:

- absent: Absence of the TE (reference allele)
- present: Presence of the TE insertion (as genotyped by Paragraph)
- excised: Presence of the 6-bp insertion putatively left by TE excision (as genotyped by Platypus)

The number of samples in each group was as follows:

- absent: 71

- present: 9
- excised: 14

All genotype calls for the TE insertion were homozygous and were therefore easy to classify as "present" when homozygous for the alternate allele. For the 6-bp insertion, however, some genotype calls were heterozygous. In all such cases, visual analysis of the aligned Illumina reads confirmed that these were most likely mis-genotyped and were actually homozygous for the alternate allele. Therefore, samples that were heterozygous at that site were classified as "excised" in addition to those that were homozygous for the alternate allele. Samples that were called as homozygous for the reference allele at the 6-bp insertion position were classified as "absent". Finally, eight (8) samples for which missing genotype calls did not allow unambiguous classification were discarded and not used for further analysis.

Using this classification, we computed the alternate allele frequency within each of the three allele classes of the 156 SNVs in the 39-kb LD block. These results are visualized in figure 5d of the main text. They show clear similarity between the "present" and "excised" haplotypes, which is indeed consistent with the 6-bp insertion being a remnant from the excision of the 480-bp TE.

**Table S1:** Number of filtered variants per SV type and size class discovered by Sniffles for 17 samples subjected to Oxford Nanopore sequencing. These SVs were used as a truth set for the benchmarks presented in this paper.

| | [50bp – 100bp[ | | | | [100bp – 1kb[ | | | | [1kb – 10kb[ | | | | ≥ 10kb | | | |
|---|---|---|---|---|---|---|---|---|---|---|---|---|---|---|---|---|
| | DEL[a] | INS[b] | DUP[c] | INV[d] | DEL | INS | DUP | INV | DEL | INS | DUP | INV | DEL | INS | DUP | INV |
| AC2001 | 2295 | 2591 | 23 | 3 | 2432 | 2643 | 541 | 56 | 1533 | 873 | 369 | 43 | 409 | 14 | 79 | 37 |
| Alta | 3389 | 3614 | 30 | 7 | 3441 | 4085 | 802 | 105 | 2321 | 1732 | 905 | 83 | 625 | 60 | 191 | 83 |
| Maple Isle | 3443 | 3546 | 19 | 7 | 3480 | 3910 | 597 | 90 | 2220 | 1492 | 496 | 69 | 533 | 46 | 147 | 70 |
| Maple Presto | 3037 | 3162 | 7 | 4 | 2904 | 2920 | 359 | 38 | 1616 | 362 | 197 | 29 | 361 | 0 | 34 | 26 |
| OAC 09-35C | 2861 | 2949 | 9 | 4 | 2887 | 3280 | 565 | 68 | 1970 | 1289 | 496 | 45 | 495 | 43 | 114 | 46 |
| OAC Carman | 2895 | 3006 | 13 | 4 | 2716 | 2830 | 425 | 30 | 1588 | 331 | 355 | 34 | 347 | 0 | 33 | 15 |
| OAC Drayton | 2473 | 2761 | 5 | 3 | 2551 | 2923 | 509 | 61 | 1686 | 1073 | 397 | 39 | 402 | 40 | 119 | 30 |
| OAC Embro | 2944 | 3176 | 6 | 4 | 2930 | 3150 | 408 | 47 | 1635 | 802 | 310 | 34 | 353 | 9 | 76 | 26 |
| OAC Lakeview | 2697 | 2993 | 16 | 4 | 2780 | 3313 | 650 | 69 | 1700 | 1122 | 602 | 54 | 384 | 32 | 125 | 46 |
| OAC Madoc | 2679 | 2945 | 10 | 2 | 2748 | 2935 | 449 | 61 | 1818 | 833 | 309 | 40 | 418 | 6 | 86 | 32 |
| OAC Oxford | 2678 | 2967 | 10 | 6 | 2772 | 3416 | 677 | 88 | 1809 | 1418 | 686 | 56 | 439 | 95 | 198 | 64 |
| OAC Petrel | 2670 | 2902 | 9 | 4 | 2746 | 3180 | 556 | 62 | 1893 | 1304 | 516 | 48 | 410 | 77 | 113 | 43 |
| OAC Prudence | 3095 | 3323 | 13 | 8 | 3249 | 3647 | 655 | 92 | 2140 | 1485 | 631 | 66 | 545 | 65 | 175 | 55 |
| OAC Stratford | 2676 | 2936 | 9 | 6 | 2794 | 3179 | 574 | 68 | 1865 | 1251 | 502 | 46 | 463 | 62 | 126 | 46 |
| OT09-03 | 2983 | 3229 | 3 | 3 | 3057 | 3387 | 529 | 58 | 2013 | 1380 | 467 | 34 | 462 | 71 | 104 | 39 |
| QS5091.50j | 2433 | 2647 | 9 | 6 | 2486 | 2525 | 375 | 45 | 1472 | 424 | 216 | 32 | 369 | 0 | 67 | 31 |
| Roland | 3001 | 3118 | 12 | 6 | 3093 | 3429 | 551 | 55 | 2075 | 1426 | 533 | 51 | 477 | 72 | 131 | 42 |

[a] DEL: deletions
[b] INS: insertions
[c] DUP: duplications
[d] INV: inversions

**Table S2:** Number of deletions and insertions overlapping various genic features and proportion of these features within the reference annotation of Williams82

| | Whole genome | | | Non-repeated genome | | |
|---|---|---|---|---|---|---|
| Feature | % in reference | DEL[a] (%) | INS[b] (%) | % in reference | DEL (%) | INS (%) |
| cds[c] | 6.4 | 1685 (5.5) | 386 (1.5) | 11.0 | 1617 (7.2) | 372 (2.4) |
| gene | 15.5 | 4329 (14.1) | 3989 (15.2) | 24.1 | 3806 (17.0) | 3161 (20.3) |
| intergenic | 60.0 | 15793 (51.5) | 13893 (53.0) | 39.3 | 9628 (43.0) | 6509 (41.7) |
| upstream5kb[d] | 18.1 | 8853 (28.9) | 7967 (30.4) | 25.6 | 7343 (32.8) | 5562 (35.6) |

[a] DEL: deletions
[b] INS: insertions
[c] cds: coding sequence
[d] upstream5kb: within 5 kb upstream of a gene

**Table S3:** Randomization test on the mean allele frequencies of SVs depending on the genic features overlapped

| | Deletions | | Insertions | |
|---|---|---|---|---|
| Genic features | Observed difference[a] | p-value[b] | Observed difference | p-value |
| cds[c]- gene | -0.0499 | $< 10^{-4}$ | 0.0015 | 0.465 |
| cds - intergenic | -0.0579 | $< 10^{-4}$ | -0.0199 | 0.085 |
| cds - upstream5kb[d] | -0.0581 | $< 10^{-4}$ | 0.0012 | 0.473 |
| gene - intergenic | -0.0081 | 0.037 | -0.0214 | $< 10^{-4}$ |
| gene - upstream5kb | -0.0082 | 0.049 | -0.0003 | 0.477 |
| intergenic - upstream5kb | -0.0001 | 0.484 | 0.0211 | $< 10^{-4}$ |

[a] Observed difference between the mean allele frequencies of the SVs overlapping the corresponding genic features

[b] p-value computed from a one-sided comparison to the distribution of mean differences obtained from the randomizations (10,000 iterations). The significance threshold was set to $\alpha = 0.05 \div 6 = 0.008$ to correct for multiple testing.

[c] cds: coding sequence

[d] upstream5kb: within 5 kb upstream of a gene

**Table S4:** List of Gene Ontology Biological Process terms that are overrepresented ($\alpha = 0.05$) among coding sequences impacted by SVs with frequency $> 0.5$

| GO ID | Description | Number of genes[a] | Expected[b] | Observed[c] | p-value[d] |
|---|---|---|---|---|---|
| GO:0006952 | defense response | 3218 | 31.86 | 73 | 2.771e-08 |
| GO:1901420 | negative regulation of response to alcohol | 157 | 1.57 | 10 | 0.006 |
| GO:0009788 | negative regulation of abscisic acid-activated signaling pathway | 157 | 1.57 | 10 | 0.006 |
| GO:0032922 | circadian regulation of gene expression | 52 | 0.52 | 6 | 0.01771 |
| GO:0010483 | pollen tube reception | 34 | 0.34 | 5 | 0.02852 |
| GO:1905957 | regulation of cellular response to alcohol | 334 | 3.33 | 13 | 0.0494 |

[a] Number of genes with corresponding GO annotation within the gene universe (whole genome except unanchored scaffolds)
[b] Expected number of genes with given GO annotation within the subset of interest considering a hypergeometric distribution
[c] Observed number of genes with given GO annotation within the subset of interest
[d] Bonferroni-corrected p-values computed using the conditional hypergeometric test in the GOstats R package [19]

**Table S5:** List of Gene Ontology Biological Process terms that are underrepresented ($\alpha = 0.05$) among coding sequences impacted by SVs with frequency $> 0.5$

| GO ID | Description | Number of genes[a] | Expected[b] | Observed[c] | p-value[d] |
|---|---|---|---|---|---|
| GO:1901576 | organic substance biosynthetic process | 10207 | 101.83 | 57 | 7.826e-05 |
| GO:0034641 | cellular nitrogen compound metabolic process | 9963 | 99.39 | 56 | 0.0001638 |
| GO:0009058 | biosynthetic process | 10423 | 103.98 | 60 | 0.000188 |
| GO:0009059 | macromolecule biosynthetic process | 7143 | 71.26 | 34 | 0.0002087 |
| GO:0044271 | cellular nitrogen compound biosynthetic process | 6818 | 68.02 | 32 | 0.0003073 |
| GO:0044237 | cellular metabolic process | 20215 | 201.67 | 151 | 0.0004943 |
| GO:0034645 | cellular macromolecule biosynthetic process | 6988 | 69.71 | 34 | 0.0005673 |
| GO:0044249 | cellular biosynthetic process | 9988 | 99.64 | 58 | 0.000584 |
| GO:0008152 | metabolic process | 23463 | 234.07 | 185 | 0.00105 |
| GO:0034654 | nucleobase-containing compound biosynthetic process | 5166 | 51.54 | 22 | 0.002042 |
| GO:1901360 | organic cyclic compound metabolic process | 9577 | 95.54 | 57 | 0.002998 |
| GO:0018130 | heterocycle biosynthetic process | 5705 | 56.91 | 27 | 0.005327 |
| GO:0006725 | cellular aromatic compound metabolic process | 9332 | 93.10 | 56 | 0.005933 |
| GO:0019438 | aromatic compound biosynthetic process | 5953 | 59.39 | 29 | 0.00599 |
| GO:0006139 | nucleobase-containing compound metabolic process | 8237 | 82.17 | 47 | 0.006126 |
| GO:0046483 | heterocycle metabolic process | 8933 | 89.12 | 53 | 0.007338 |
| GO:0016070 | RNA metabolic process | 6800 | 67.84 | 36 | 0.008725 |
| GO:1901362 | organic cyclic compound biosynthetic process | 6214 | 61.99 | 32 | 0.01332 |
| GO:0044238 | primary metabolic process | 19412 | 193.66 | 150 | 0.01586 |
| GO:0032774 | RNA biosynthetic process | 4661 | 46.50 | 21 | 0.02269 |
| GO:0097659 | nucleic acid-templated transcription | 4648 | 46.37 | 21 | 0.02466 |
| GO:0071704 | organic substance metabolic process | 20977 | 209.27 | 167 | 0.03247 |
| GO:0010556 | regulation of macromolecule biosynthetic process | 4874 | 48.62 | 23 | 0.03312 |
| GO:0051171 | regulation of nitrogen compound metabolic process | 5638 | 56.25 | 29 | 0.03994 |
| GO:2000112 | regulation of cellular macromolecule biosynthetic process | 4842 | 48.30 | 23 | 0.04037 |
| GO:0090304 | nucleic acid metabolic process | 7519 | 75.01 | 44 | 0.04127 |
| GO:0051252 | regulation of RNA metabolic process | 4681 | 46.70 | 22 | 0.04733 |
| GO:0031323 | regulation of cellular metabolic process | 6248 | 62.33 | 34 | 0.04791 |

[a] Number of genes with corresponding GO annotation within the gene universe (whole genome except unanchored scaffolds)
[b] Expected number of genes with given GO annotation within the subset of interest considering a hypergeometric distribution
[c] Observed number of genes with given GO annotation within the subset of interest
[d] Bonferroni-corrected p-values computed using the conditional hypergeometric test in the GOstats R package [19]

**Table S6:** List of PFAM domains that are overrepresented ($\alpha = 0.05$) among coding sequences impacted by SVs with frequency $> 0.5$

| PFAM ID | Description | Number of genes[a] | Expected[b] | Observed[c] | p-value[d] |
|---|---|---|---|---|---|
| PF00931 | NB-ARC domain | 353 | 3.46 | 42 | 3.011e-30 |
| PF13676 | TIR domain | 156 | 1.53 | 20 | 1.844e-14 |
| PF12776 | Myb/SANT-like DNA-binding domain | 90 | 0.88 | 14 | 4.998e-11 |
| PF05699 | hAT family C-terminal dimerisation region | 98 | 0.96 | 10 | 8.656e-06 |
| PF13947 | Wall-associated receptor kinase galacturonan-binding | 76 | 0.74 | 8 | 0.0001527 |
| PF14380 | Wall-associated receptor kinase C-terminal | 36 | 0.35 | 6 | 0.0002372 |
| PF13855 | Leucine rich repeat | 603 | 5.90 | 20 | 0.0004999 |
| PF08263 | Leucine rich repeat N-terminal domain | 555 | 5.43 | 19 | 0.000564 |
| PF14291 | Domain of unknown function (DUF4371) | 44 | 0.43 | 6 | 0.0008046 |
| PF00560 | Leucine Rich Repeat | 443 | 4.34 | 16 | 0.001755 |
| PF07714 | Protein tyrosine and serine/threonine kinase | 1089 | 10.66 | 25 | 0.01602 |
| PF13962 | Domain of unknown function | 87 | 0.85 | 6 | 0.04042 |

[a] Number of genes with corresponding PFAM annotation within the gene universe (whole genome except unanchored scaffolds)
[b] Expected number of genes with given PFAM annotation within the subset of interest considering a hypergeometric distribution
[c] Observed number of genes with given PFAM annotation within the subset of interest
[d] Bonferroni-corrected p-values computed using the hypergeometric test in the GOstats R package [19]

**Table S7:** DNA extraction and preparation methods for Oxford Nanopore sequencing

| Sample[a] | Growth[b] | Grinding[c] | Extraction[d] | RNase T (°C)[e] | Size selection[f] | Combined[g] |
|---|---|---|---|---|---|---|
| AC2001 (1) | Lab | LN + TL | CTAB | 37 | BP 15kb | no |
| AC2001 (2) | Lab | LN + TL | CTAB | 37 | SRE | no |
| Alta | Lab | LN + TL | CTAB | 37 | BP 15kb | no |
| Maple Isle (1) | Lab | LN + TL | CTAB | 37 | BP 15kb | no |
| Maple Isle (2) | Lab | LN + TL | CTAB | 60 | SRE | no |
| Maple Presto | Field | CD + TL | DNeasy | 65 | BP 6kb | yes |
| OAC 09-35C | Lab | LN + TL | CTAB | 37 | BP 15kb | no |
| OAC Carman | Field | CD + TL | DNeasy | 65 | BP 6kb | yes |
| OAC Drayton | Lab | LN + TL | CTAB | 37 | BP 15kb | no |
| OAC Embro | Greenhouse | LN + MP | Gentra | 37 | None | no |
| OAC Lakeview (1) | Lab | LN + TL | CTAB | 37 | BP 15kb | no |
| OAC Lakeview (2) | Lab | LN + TL | CTAB | 60 | SRE | no |
| OAC Madoc | Lab | LN + TL | CTAB | 37 | BP 15kb | no |
| OAC Oxford | Lab | LN + TL | CTAB | 37 | BP 15kb | no |
| OAC Petrel | Lab | LN + TL | CTAB | 60 | SRE | no |
| OAC Prudence | Lab | LN + TL | CTAB | 37 | BP 15kb | no |
| OAC Stratford | Lab | LN + TL | CTAB | 37 | BP 15kb | no |
| OT09-03 | Lab | LN + TL | CTAB | 65 | SRE | no |
| QS5091.50j | Field | CD + TL | DNeasy | 65 | BP 6kb | no |
| Roland | Lab | LN + TL | CTAB | 60 | SRE | no |

[a] The number in parentheses identifies samples sequenced on two different flowcells

[b] Field: leaf harvested at Agriculture and Agri-Food Canada in Ottawa, and cryodessicated on site

[c] Grinding method: LN + TL, Frozen in liquid nitrogen and ground using Qiagen TissueLyser;
CD + TL, Cryodessicated at harvest and ground using Qiagen TissueLyser;
LN + MP, Frozen in liquid nitrogen and ground using mortar and pestle

[d] Extraction protocol used: CTAB, CTAB-based protocol described in main text;
DNeasy, Qiagen DNeasy Plant Mini Kit;
Gentra, Qiagen Gentra Puregene Tissue Kit

[e] Incubation temperature of the RNase A treatment

[f] Size-selection protocol: None, no size-selection;
BP 6kb, BluePippin with 6-kb size-selection threshold;
BP 15kb, BluePippin High-Pass Plus cassette with 15-kb size-selection threshold;
SRE, Circulomics Short Read Eliminator Kit

[g] Indicates whether the DNA from two different extractions was combined for sequencing

**Table S8:** Metadata on Oxford Nanopore sequencing runs

| Sample[a] | Date[b] | Active pores[c] | Mass (ng)[d] | Run time (h)[e] | Yield (Gb)[f] | Read N50 (kb)[g] |
|---|---|---|---|---|---|---|
| AC2001 (1) | 13-03-2020 | 1652 | 210 | 72 | 9.3 | 6.0 |
| AC2001 (2) | 27-07-2020 | 1586 | 309 | 72 | 13.9 | 19.7 |
| Alta | 13-03-2020 | 1640 | 196 | 72 | 30.4 | 11.2 |
| Maple Isle (1) | 11-06-2020 | 1477 | 166 | 72 | 6.6 | 4.1 |
| Maple Isle (2) | 10-08-2020 | 1518 | 354 | 72 | 15.7 | 14.8 |
| Maple Presto | 01-10-2019 | 1568 | 364 | 48 | 10.2 | 5.2 |
| OAC 09-35C | 21-05-2020 | 1613 | 109 | 72 | 12.8 | 17.5 |
| OAC Carman | 09-10-2019 | 1678 | 146 | 48 | 15.6 | 3.8 |
| OAC Drayton | 15-06-2020 | 1461 | 118 | 72 | 11.1 | 18.3 |
| OAC Embro | 14-03-2019 | NA | 244 | 48 | 16.3 | 6.9 |
| OAC Lakeview (1) | 15-06-2020 | 1478 | 112 | 72 | 4.8 | 21.0 |
| OAC Lakeview (2) | 10-08-2020 | 1347 | 339 | 72 | 15.3 | 8.1 |
| OAC Madoc | 21-05-2020 | 1589 | 187 | 72 | 10.2 | 12.0 |
| OAC Oxford | 26-05-2020 | 1471 | 218 | 72 | 16.7 | 22.0 |
| OAC Petrel | 31-07-2020 | 1559 | 407 | 72 | 9.6 | 29.5 |
| OAC Prudence | 11-06-2020 | 1402 | 262 | 72 | 13.5 | 20.7 |
| OAC Stratford | 26-05-2020 | 1177 | 225 | 72 | 12.5 | 18.9 |
| OT09-03 | 25-02-2020 | 1392 | 248 | 72 | 10.1 | 24.8 |
| QS5091.50j | 11-09-2019 | 1190 | 225 | 48 | 11.4 | 6.7 |
| Roland | 07-07-2020 | 1774 | 168 | 72 | 10.9 | 25.6 |

[a] The number in parentheses identifies samples sequenced on two different flowcells
[b] Date when the sequencing run was started, format dd-mm-yyyy
[c] Number of active pores on the flowcell prior to sequencing
[d] Total estimated mass of the DNA library that was loaded on the flowcell
[e] Duration of the sequencing run
[f] Total number of bases in the fastq files following basecalling
[g] Read N50 of all reads following basecalling

**Table S9:** Classification of deletions and insertions for four samples used to test the SV refinement pipeline

| Sample | Refined[a] | Below thresholds[b] | Above max. SVLEN[c] | No alignment[d] |
|---|---|---|---|---|
| Maple Presto | 9137 | 1597 | 19 | 2881 |
| OAC Carman | 6937 | 1434 | 6 | 4192 |
| OAC Embro | 11469 | 2684 | 24 | 584 |
| QS5091.50j | 9344 | 1804 | 17 | 734 |

[a] SVs that passed all filters and had their breakpoints and/or sequence content updated by the pipeline

[b] Did not pass one or several thresholds applied when comparing the proposed refined SV to the raw SV

[c] SV length above the maximum permissible length (50 kb used here)

[d] No assembly or no alignment produced by the pipeline

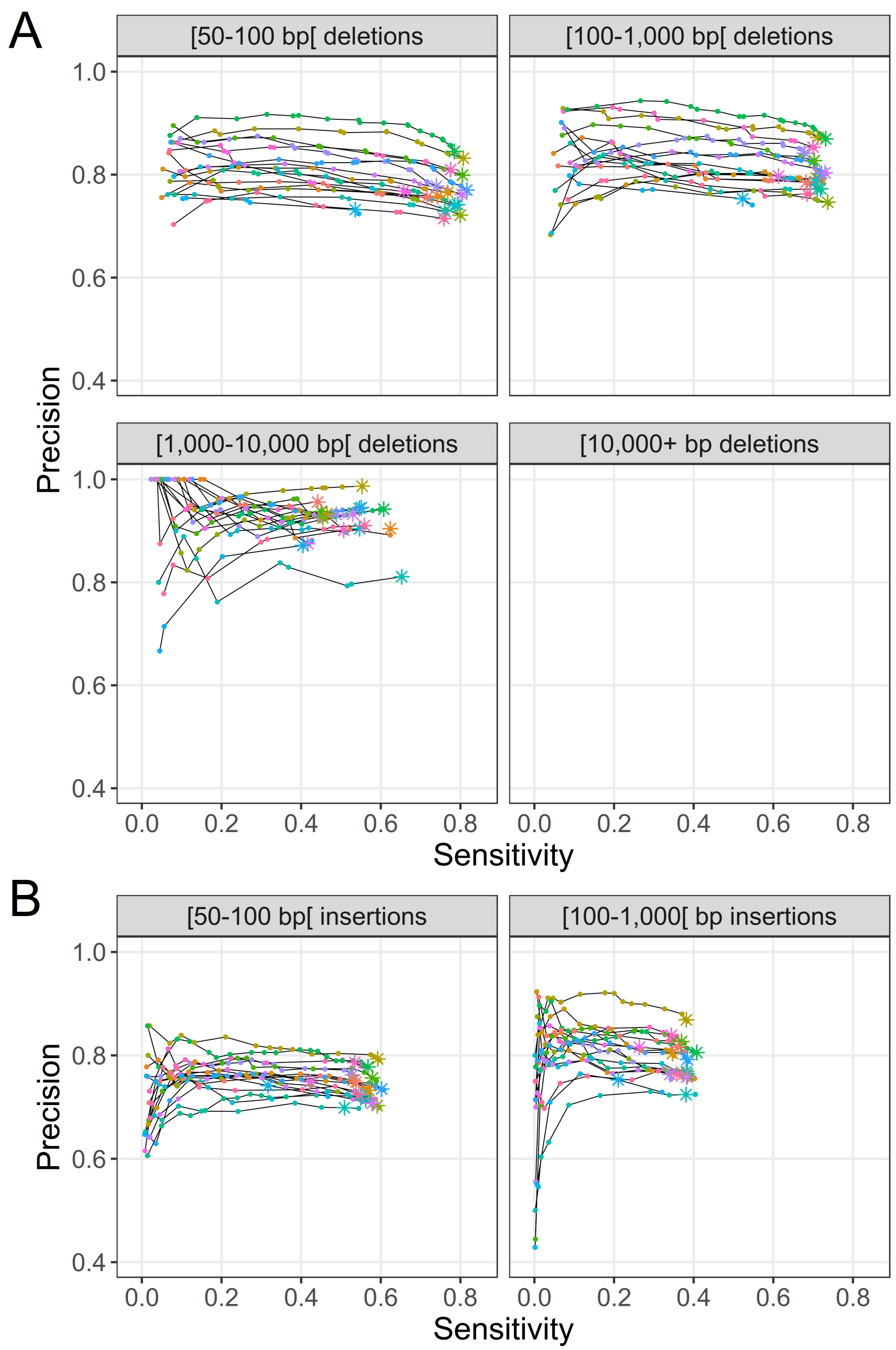

**Figure S1:** Genotyping sensitivity and precision of (A) deletions and (B) insertions discovered from the Illumina data in non-repeat regions. Only SVs with less than 20% overlap to regions annotated as repeats were used for these benchmarks. Each line and color represents one of 17 samples. The points correspond to different filtering thresholds on the minimum number of Illumina reads required to support a genotype call. The asterisks indicate a minimum number of supporting reads of 2; points to the left of these for a given line represent increasingly stringent filtering threshold values. Note that no results were available for deletions larger than 10 kb because these all overlapped repeat regions.

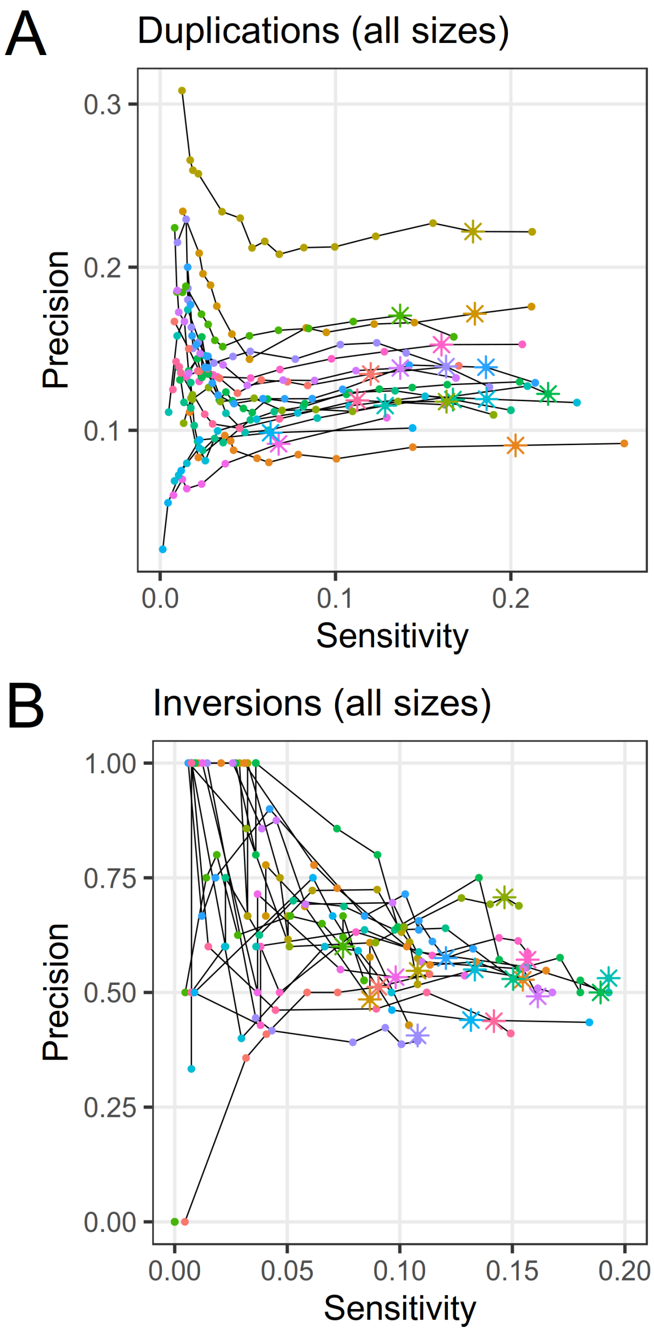

**Figure S2:** Genotyping sensitivity and precision of (A) duplications and (B) inversions discovered from the Illumina data. Each line and color represents one of 17 samples. The points correspond to different filtering thresholds on the minimum number of Illumina reads required to support a genotype call. The asterisks indicate a minimum number of supporting reads of 2; points to the left of these for a given line represent increasingly stringent filtering threshold values.

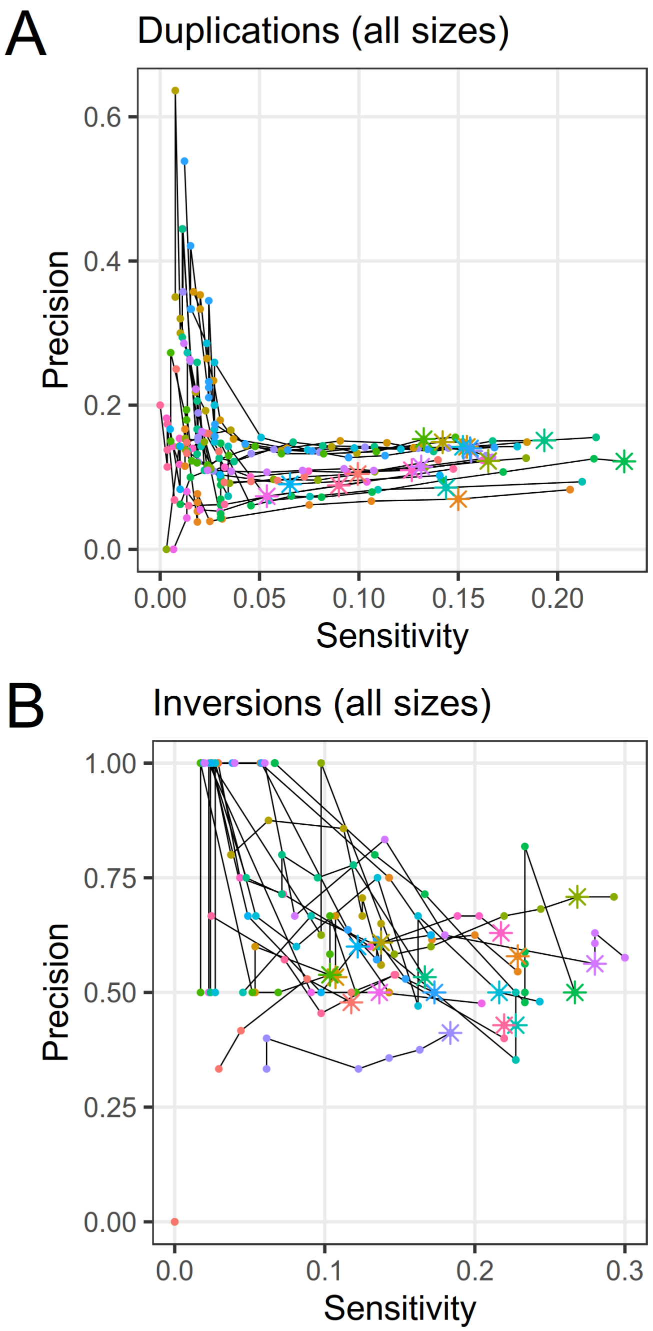

**Figure S3:** Genotyping sensitivity and precision of (A) duplications and (B) inversions discovered from the Illumina data in non-repeat regions. Only SVs with less than 20% overlap to regions annotated as repeats were used for these benchmarks. Each line and color represents one of 17 samples. The points correspond to different filtering thresholds on the minimum number of Illumina reads required to support a genotype call. The asterisks indicate a minimum number of supporting reads of 2; points to the left of these for a given line represent increasingly stringent filtering threshold values.

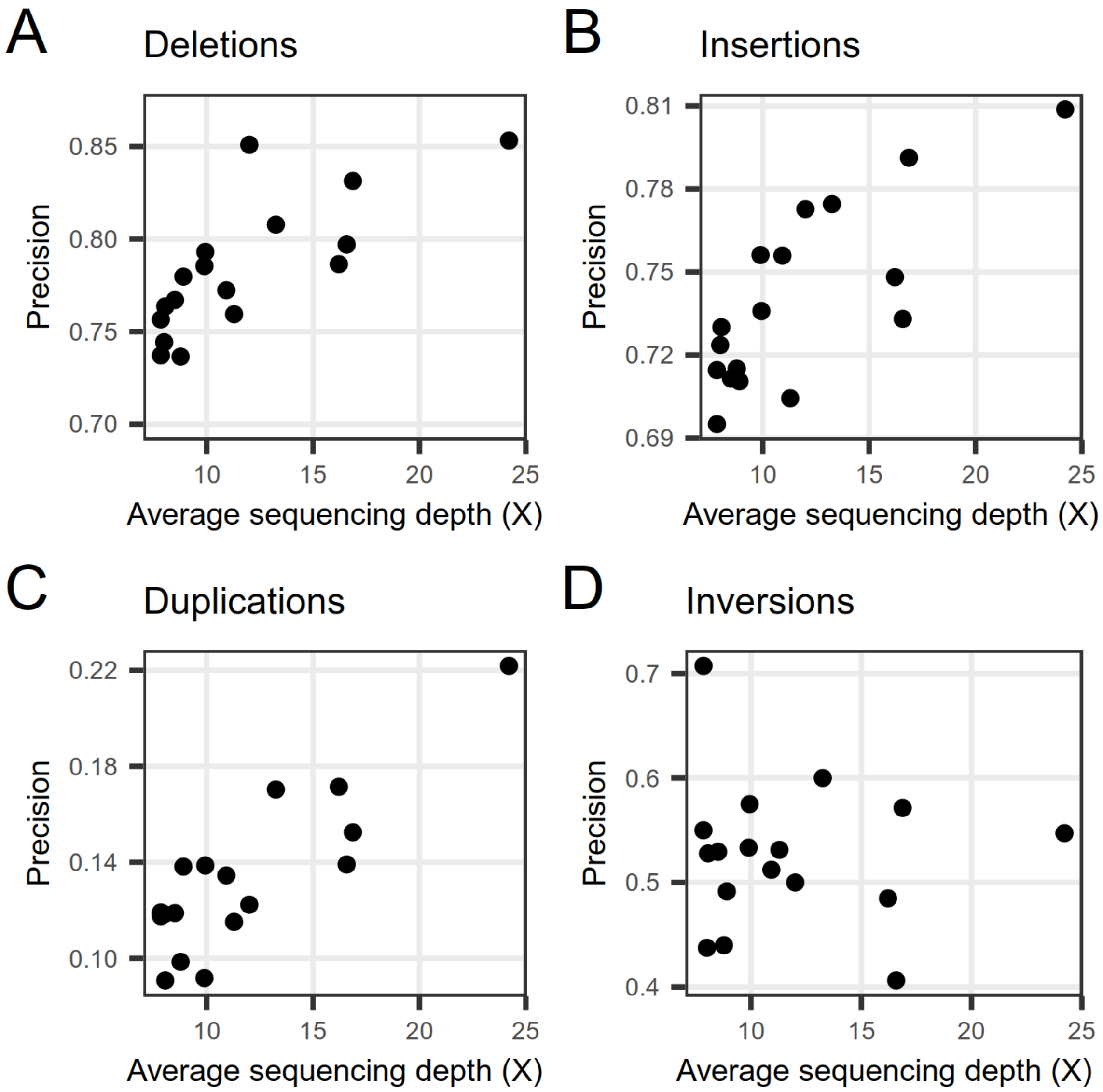

**Figure S4:** Genotyping precision of (A) deletions, (B) insertions, (C) duplications and (D) inversions discovered from Illumina data (at a minimum threshold of 2 supporting reads) for 17 samples as a function of the average Oxford Nanopore sequencing depth of that sample.

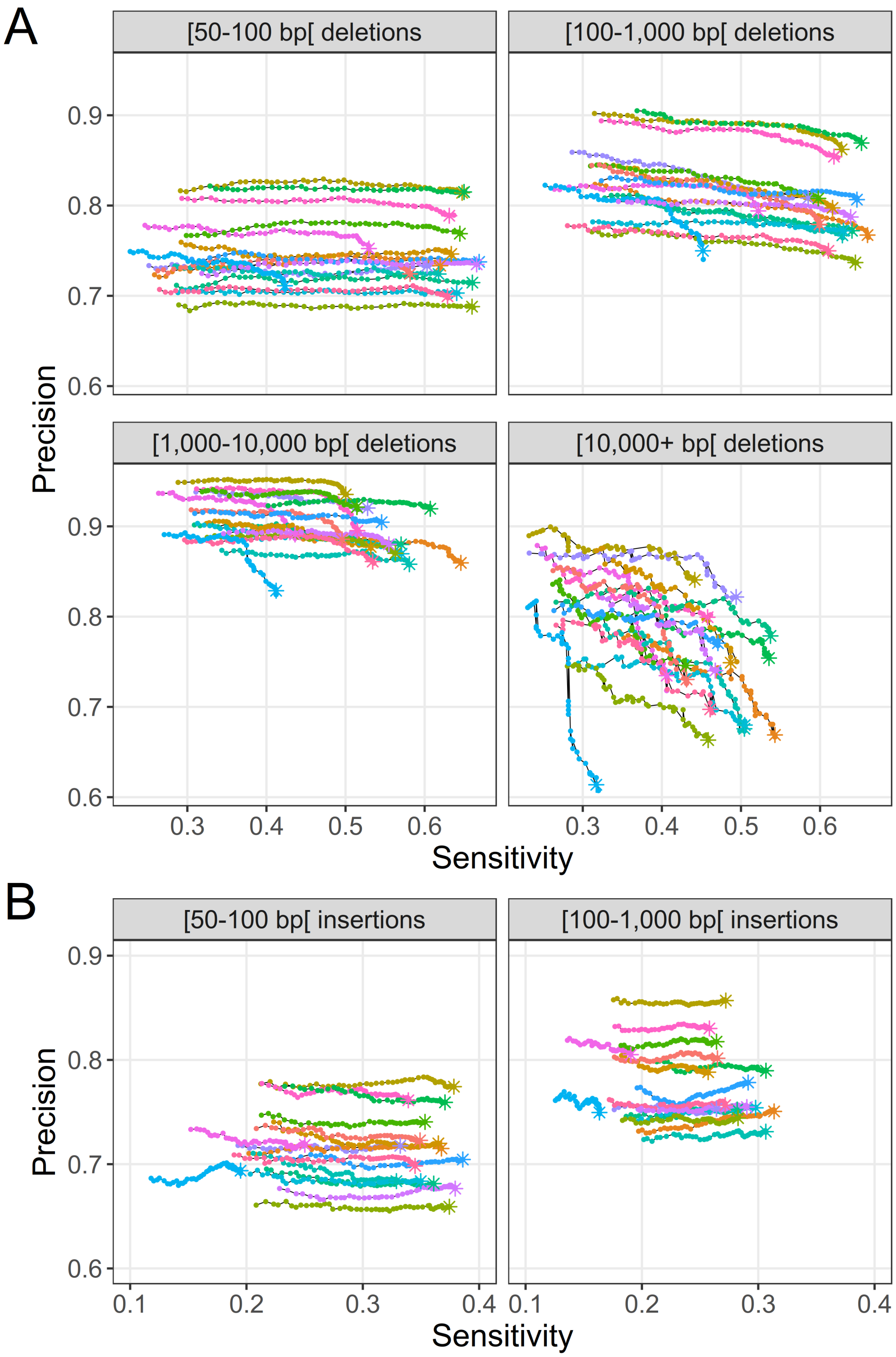

**Figure S5:** Effect of filtering variants for minimum number of ALT alleles in homozygous genotype calls (homozygous ALT count) on the genotyping performance of (A) deletions and (B) insertions discovered from Illumina data. Each line and color represents a different sample. The curve for a given sample was obtained by varying the filtering threshold for a variant to be considered for benchmarking. The asterisks mark a homozygous ALT count threshold of 4; points to the left of these for a given line represent increasingly stringent filtering threshold values.

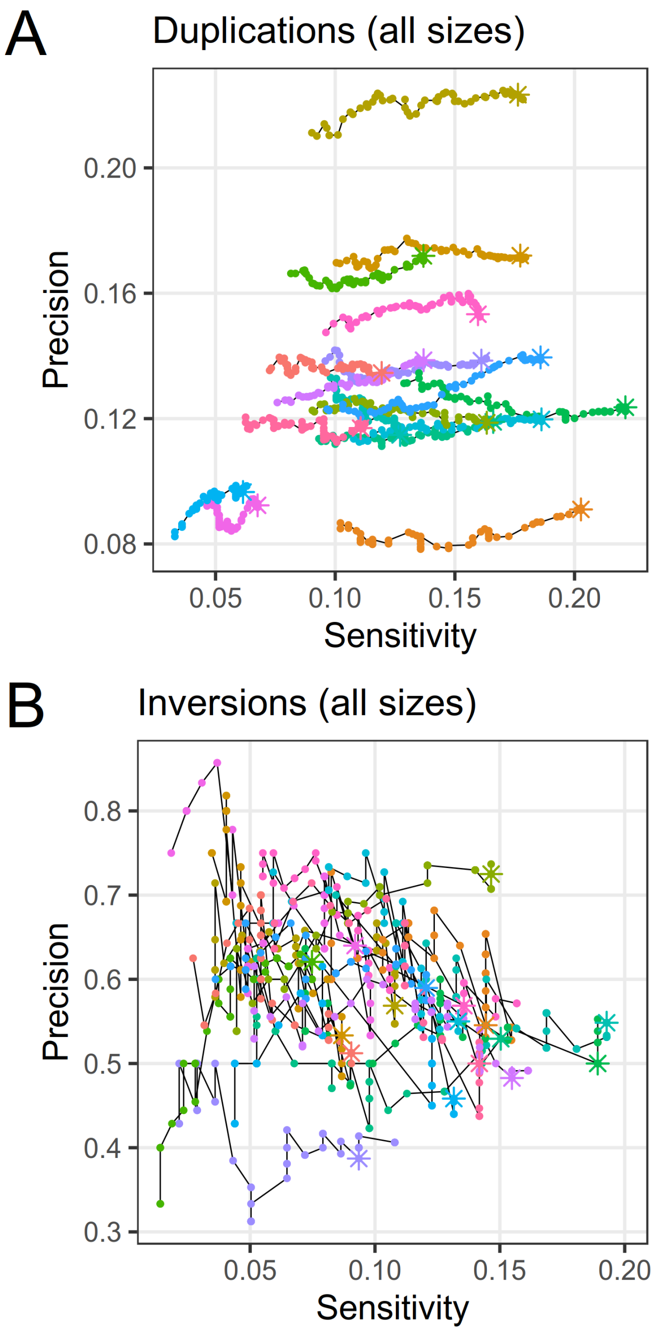

**Figure S6:** Effect of filtering variants for minimum number of ALT alleles in homozygous genotype calls (homozygous ALT count) on the genotyping performance of (A) duplications and (B) inversions discovered from Illumina data. Each line and color represents a different sample. The curve for a given sample was obtained by varying the filtering threshold for a variant to be considered for benchmarking. The asterisks mark a homozygous ALT count threshold of 6 (for duplications) or 10 (for inversions); points to the left of these for a given line represent increasingly stringent filtering threshold values.

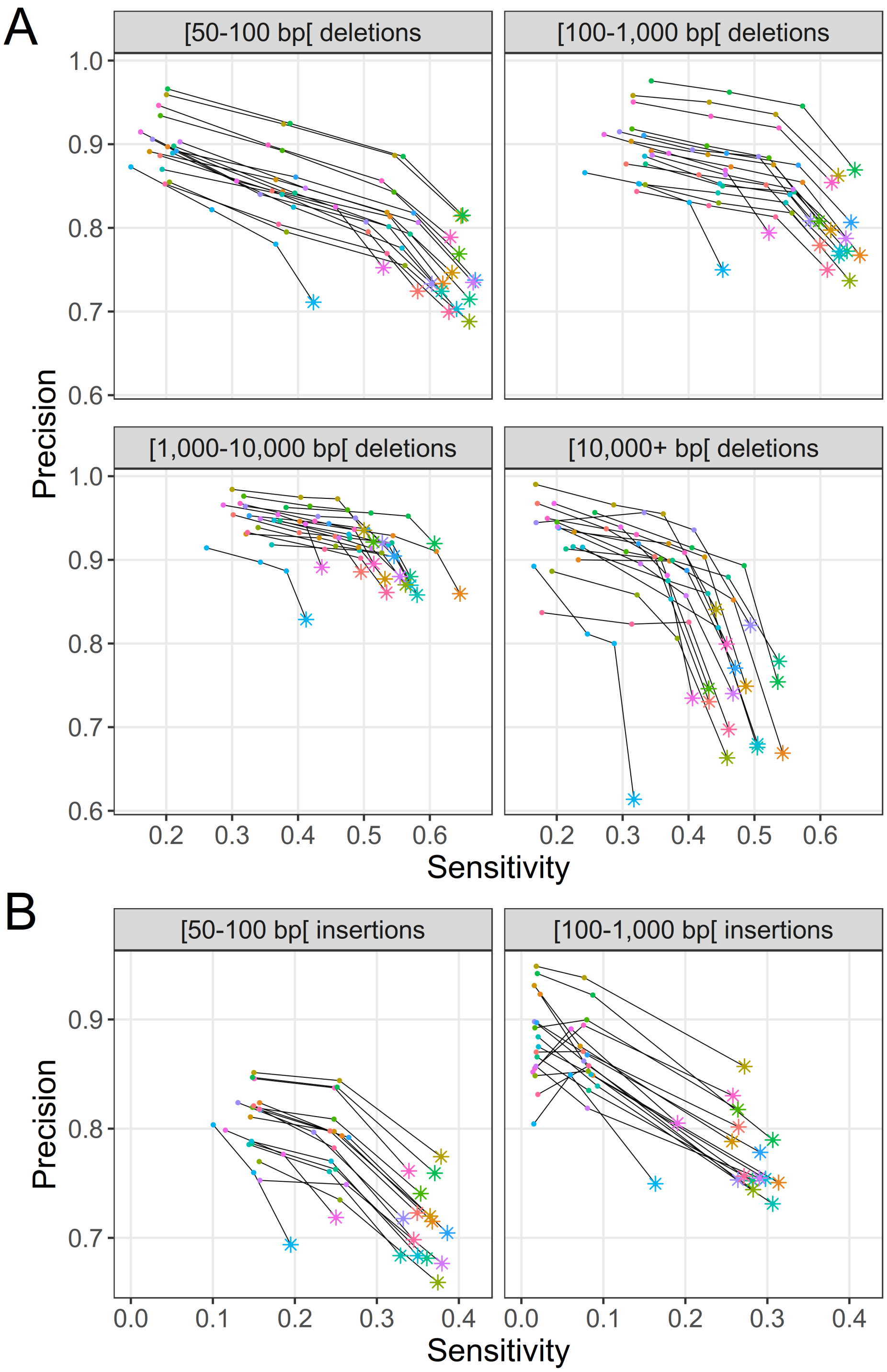

**Figure S7:** Effect of filtering variants for minimum number of supporting calling tools on the genotyping performance of (A) deletions and (B) insertions discovered from Illumina data. Each line and color represents a different sample. The curve for a given sample was obtained by varying the minimum threshold of number of supporting calling tools for a variant to be considered for benchmarking. The asterisks mark a minimum threshold of one supporting calling tool for a variant to be considered; points to the left of these for a given line represent increasingly stringent filtering threshold values.

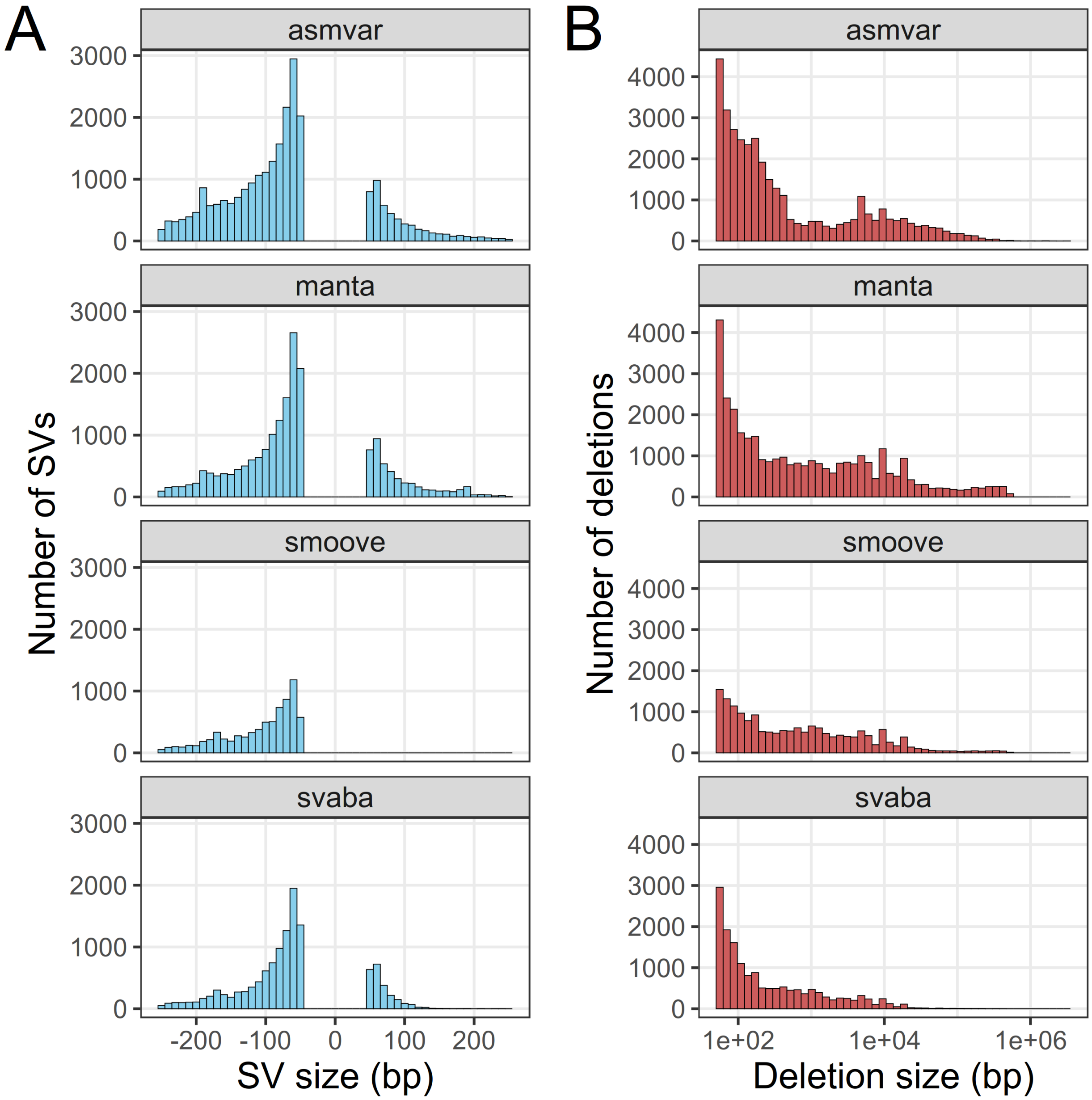

**Figure S8:** Size distribution of the SVs reported by various calling tools using Illumina data. (A) Distribution of SV sizes in the range from -250 bp to 250 bp. Negative sizes represent deletions and positive sizes represent insertions. Duplications and inversions were omitted from this plot. (B) Distribution of the absolute size of deletions on a logarithmic scale.

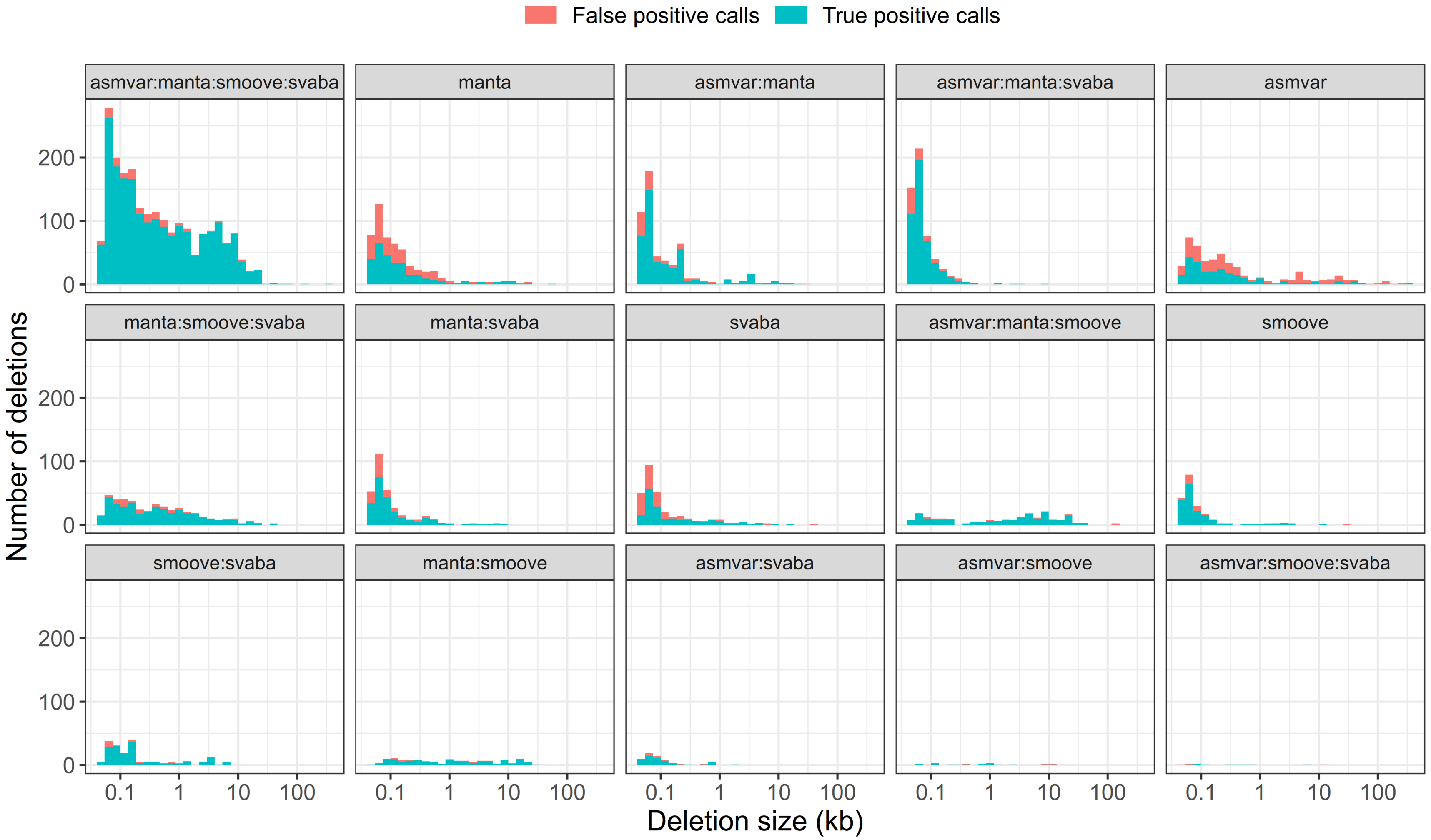

**Figure S9:** Number of true positive and false positive deletion genotype calls broken down by size and by the combination of tools that reported the variant for sample OAC Oxford. OAC Oxford was chosen as a representative sample because of its median F1 score among all 17 benchmarked samples. The genotype calls considered here were filtered for a minimum number of supporting reads of 2 and a minimum homozygous ALT count of 4 across all 102 samples in the population. Note the logarithmic scale on the x-axis.

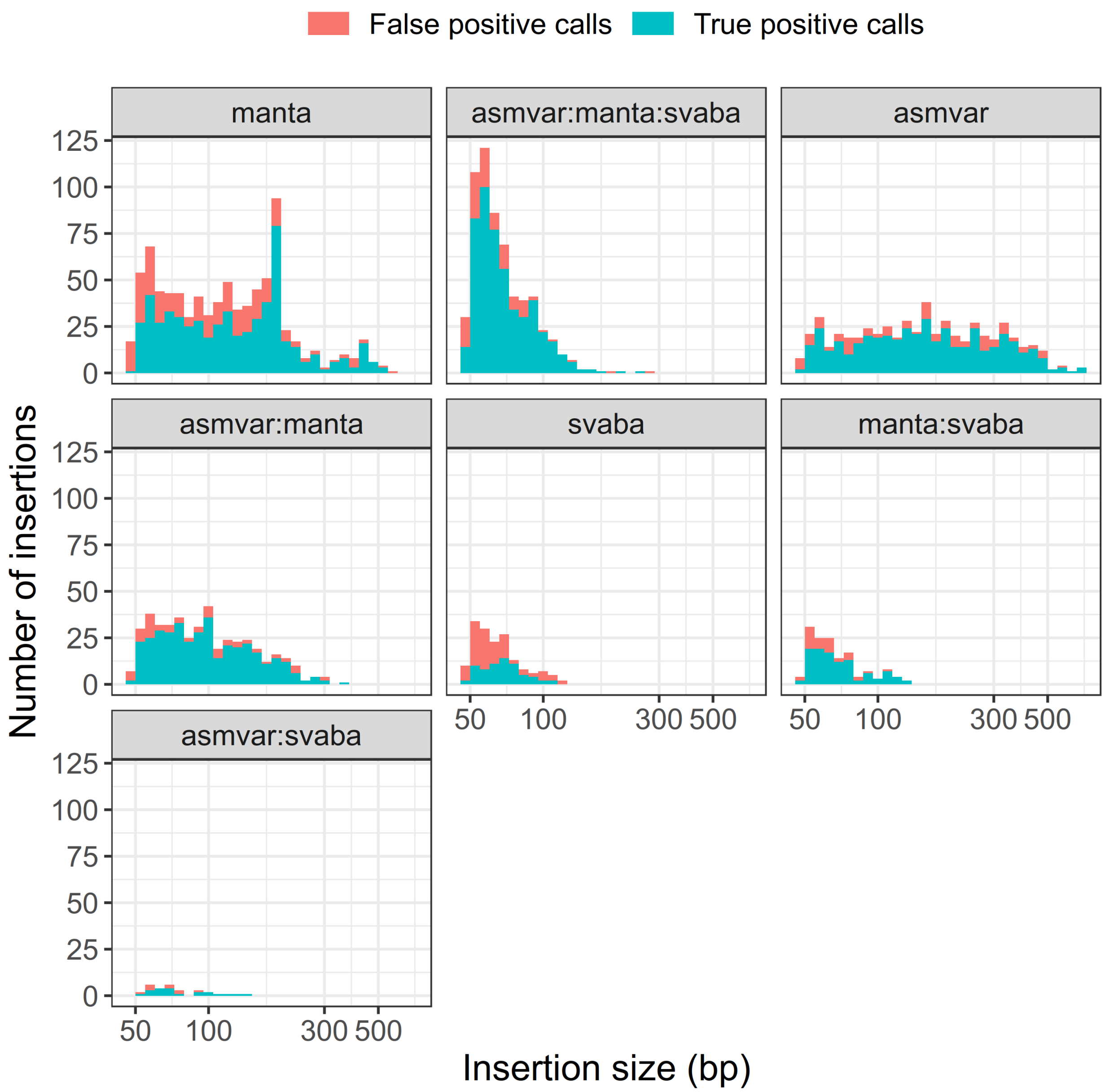

**Figure S10:** Number of true positive and false positive insertion genotype calls broken down by size and by the combination of tools that reported the variant for sample OAC Lakeview. OAC Lakeview was chosen as a representative sample because of its median F1 score among all 17 benchmarked samples. The genotype calls considered here were filtered for a minimum number of supporting reads of 2 and a minimum homozygous ALT count of 4 across all 102 samples in the population. Note the logarithmic scale on the x-axis.

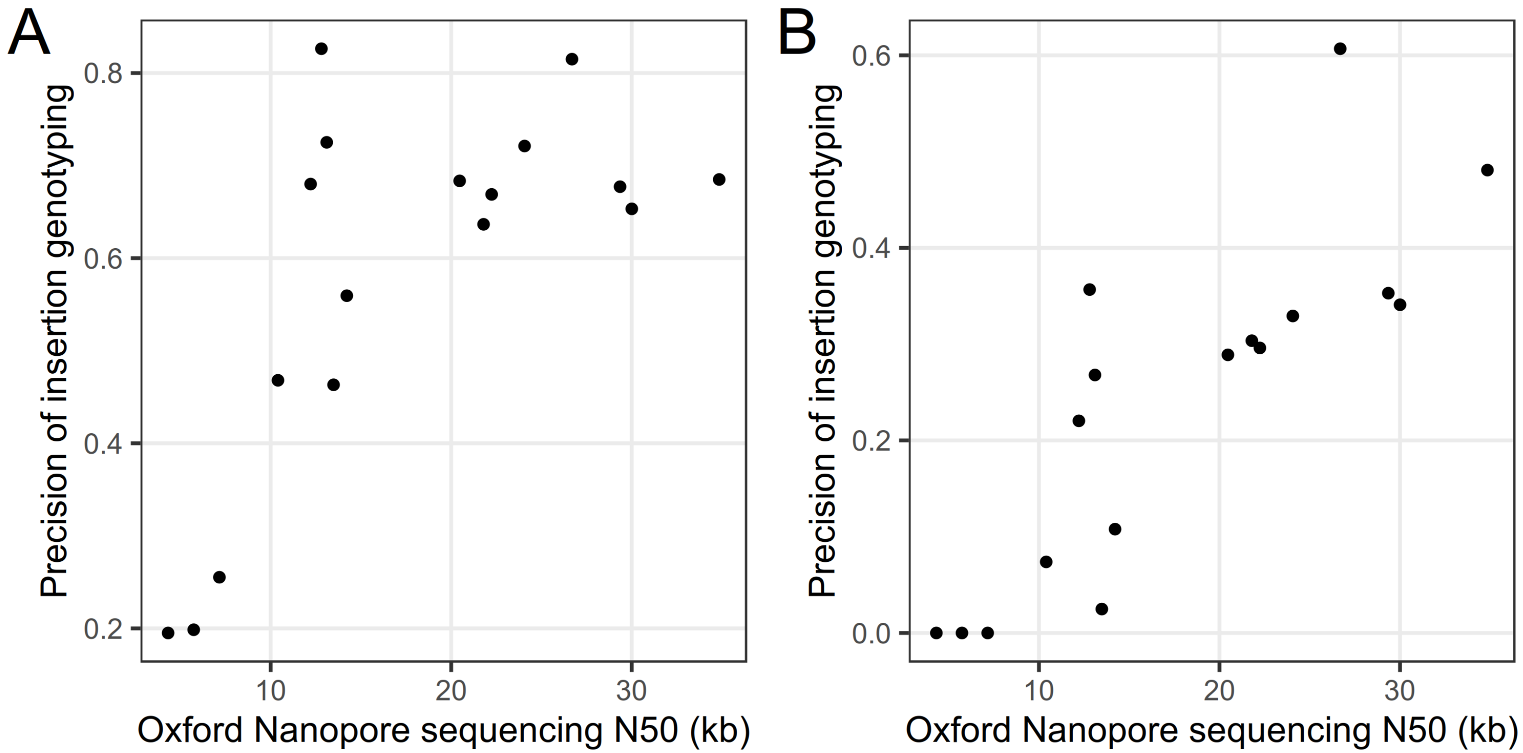

**Figure S11:** Genotyping precision of insertions discovered from Oxford Nanopore data (at a minimum threshold of 2 supporting reads) of 17 samples as a function of the read N50 of the Oxford Nanopore run(s) of that sample. (A) Precision of genotyping for insertions in the range 1,000 – 10,000 bp. (B) Precision of genotyping for insertions $\geq$ 10,000 bp.

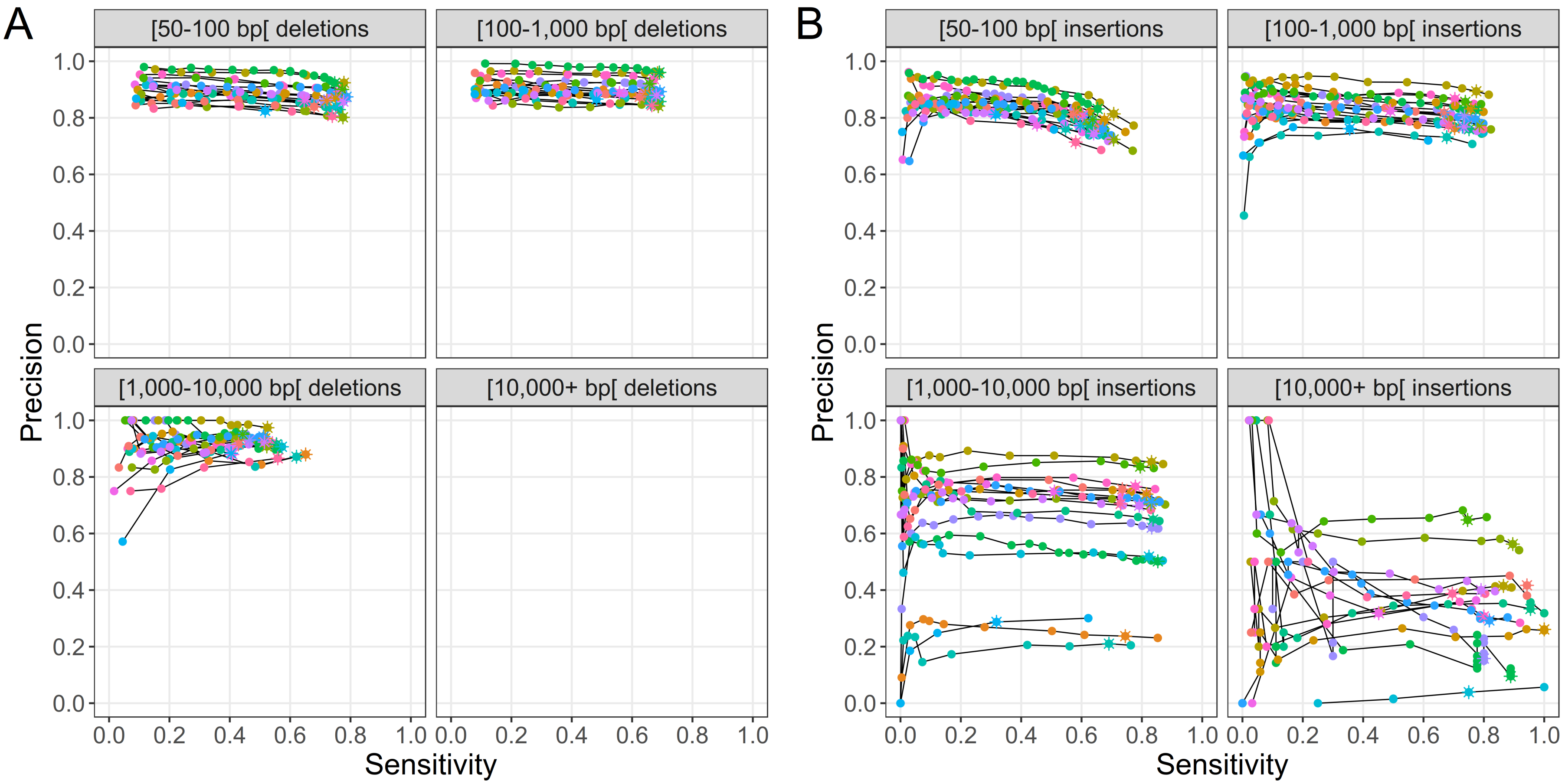

**Figure S12:** Genotyping sensitivity and precision of (A) deletions and (B) insertions discovered from the Oxford Nanopore data in non-repeat regions. Only SVs with less than 20% overlap to regions annotated as repeats were used for these benchmarks. Each line and color represents one of 17 samples. The points correspond to different filtering thresholds on the minimum number of Illumina reads required to support a genotype call. The asterisks indicate a minimum number of supporting reads of 2; points to the left of these for a given line represent increasingly stringent filtering threshold values. Note that no results were available for deletions larger than 10 kb because these all overlapped repeat regions.

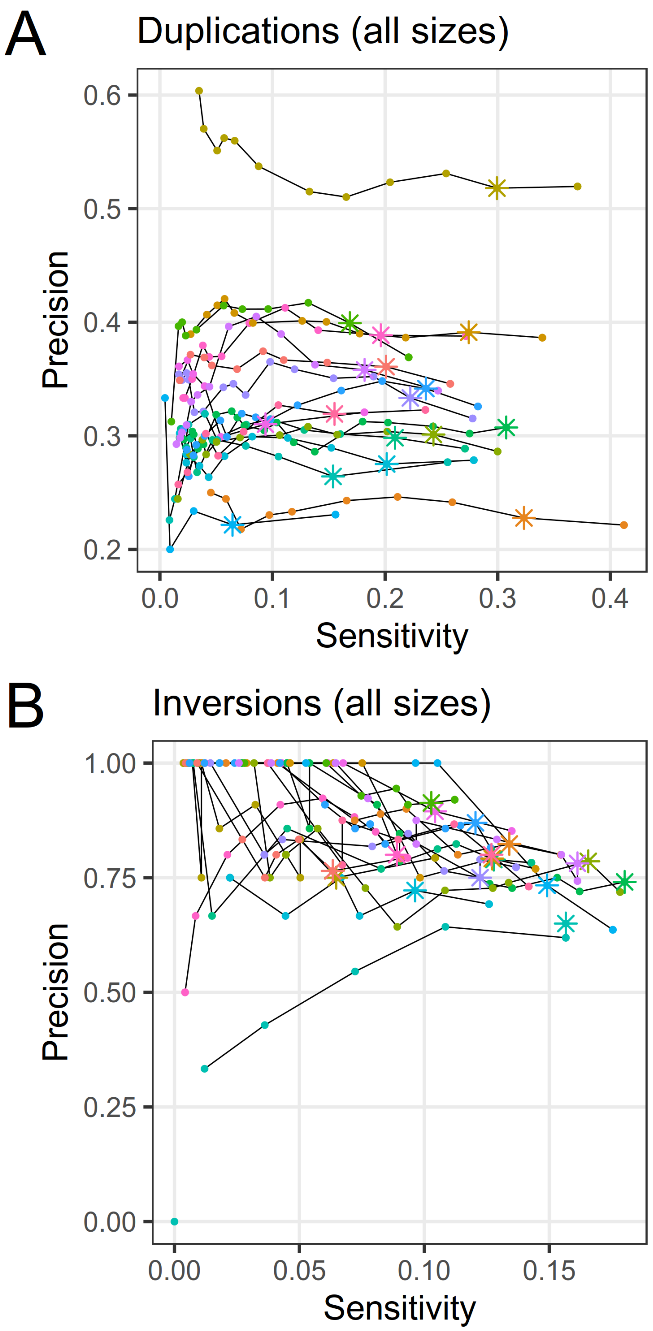

**Figure S13:** Genotyping sensitivity and precision of (A) duplications and (B) inversions discovered from the Oxford Nanopore data. Each line and color represents one of 17 samples. The points correspond to different filtering thresholds on the minimum number of Illumina reads required to support a genotype call. The asterisks indicate a minimum number of supporting reads of 2; points to the left of these for a given line represent increasingly stringent filtering threshold values.

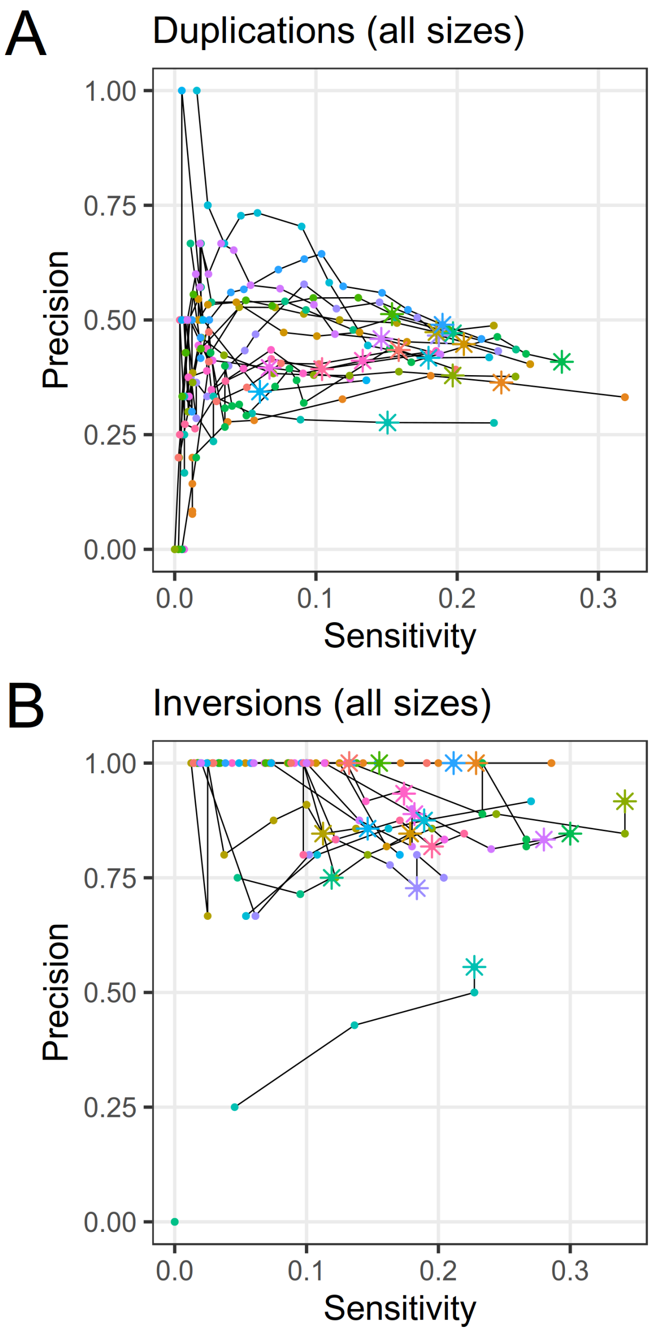

**Figure S14:** Genotyping sensitivity and precision of (A) duplications and (B) inversions discovered from the Oxford Nanopore data in non-repeat regions. Only SVs with less than 20% overlap to regions annotated as repeats were used for these benchmarks. Each line and color represents one of 17 samples. The points correspond to different filtering thresholds on the minimum number of Illumina reads required to support a genotype call. The asterisks indicate a minimum number of supporting reads of 2; points to the left of these for a given line represent increasingly stringent filtering threshold values.

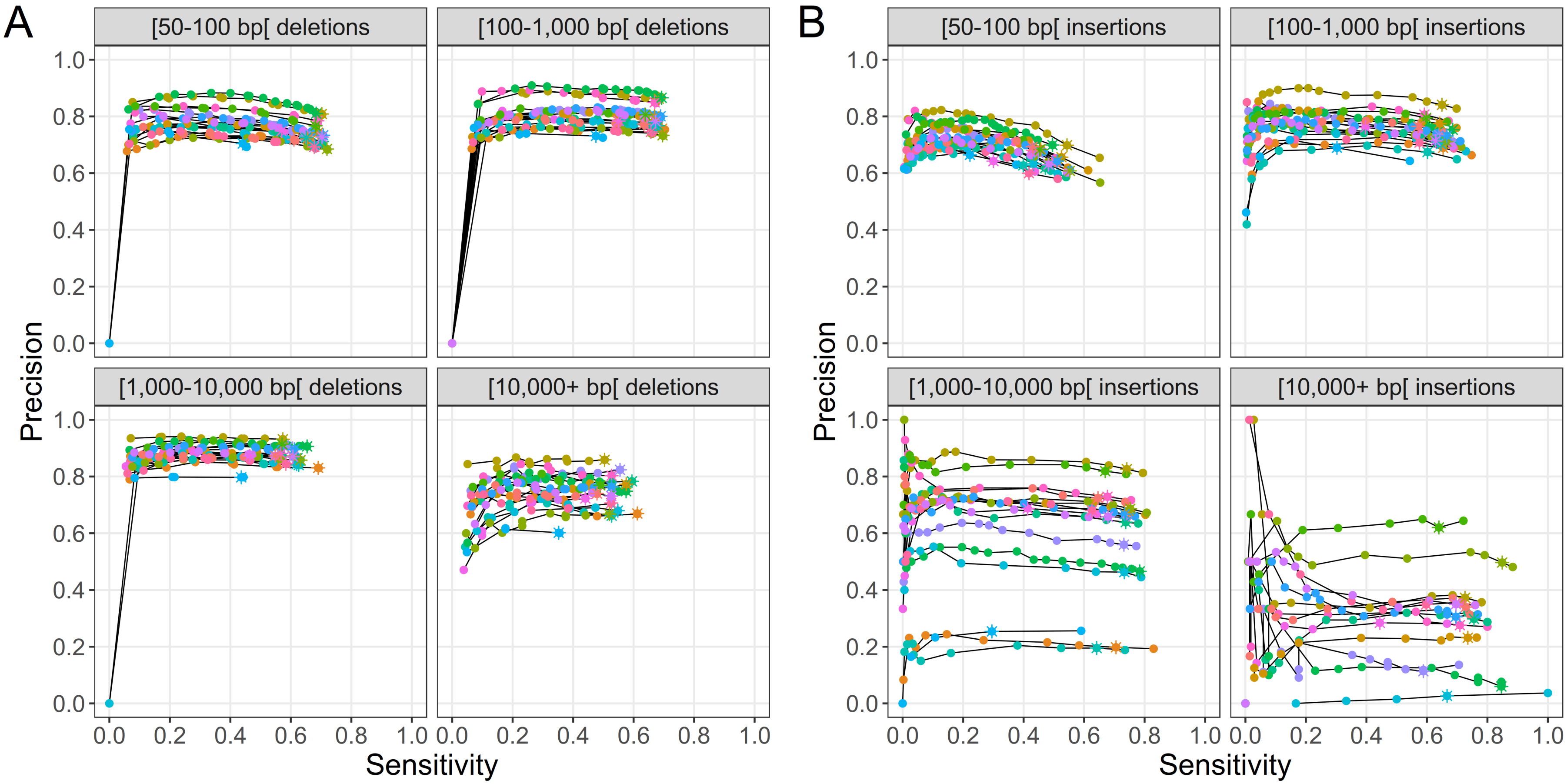

**Figure S15:** Genotyping sensitivity and precision of (A) deletions and (B) insertions in the merged dataset of SVs discovered from the Illumina and Oxford Nanopore data. Each line and color represents one of 17 samples. The points correspond to different filtering thresholds on the minimum number of Illumina reads required to support a genotype call. The asterisks indicate a minimum number of supporting reads of 2; points to the left of these for a given line represent increasingly stringent filtering threshold values.

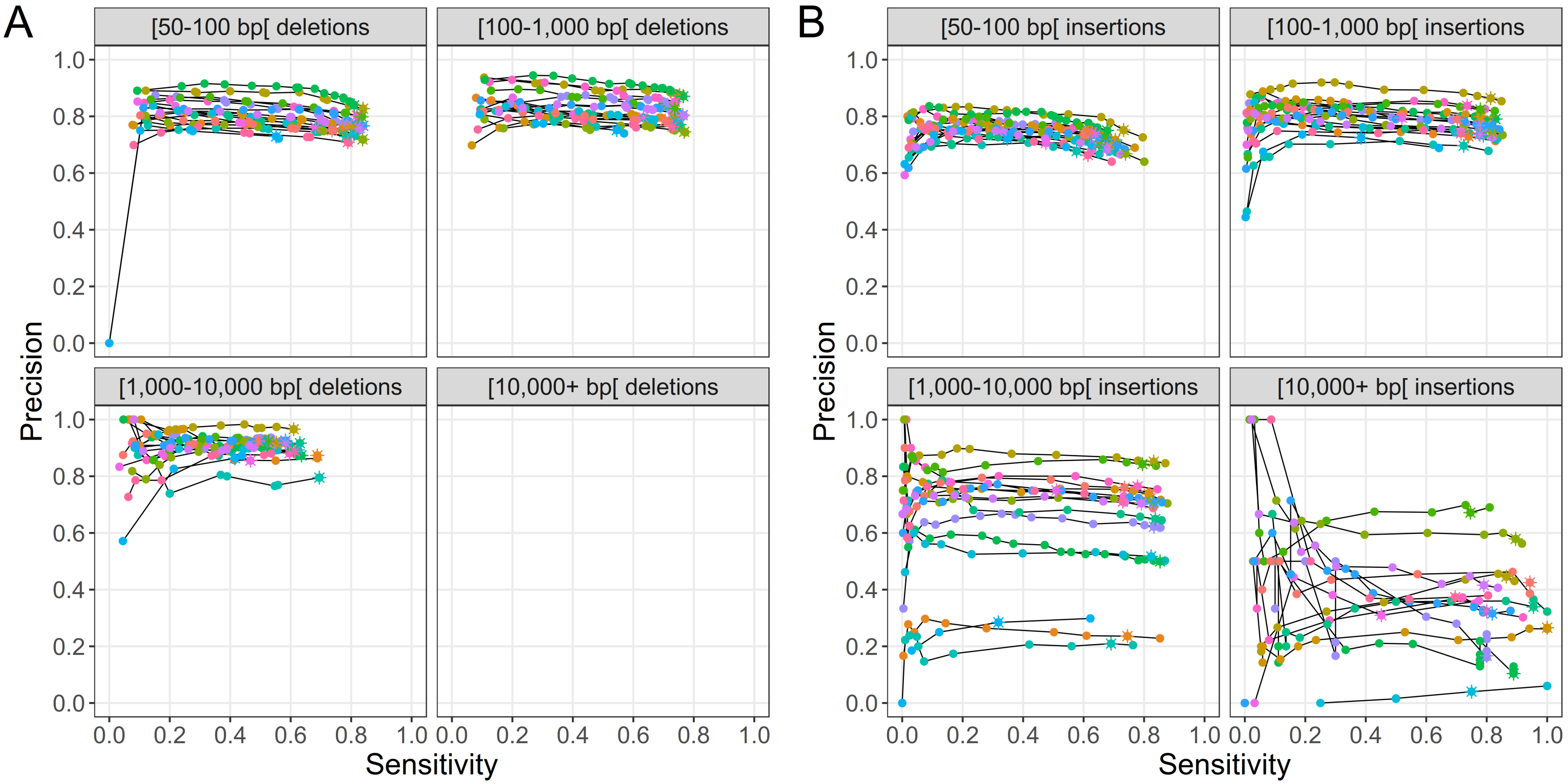

**Figure S16:** Genotyping sensitivity and precision of (A) deletions and (B) insertions in non-repeat regions in the merged dataset of SVs discovered from the Illumina and Oxford Nanopore data. Only SVs with less than 20% overlap to regions annotated as repeats were used for these benchmarks. Each line and color represents one of 17 samples. The points correspond to different filtering thresholds on the minimum number of Illumina reads required to support a genotype call. The asterisks indicate a minimum number of supporting reads of 2; points to the left of these for a given line represent increasingly stringent filtering threshold values. Note that no results were available for deletions larger than 10 kb because these all overlapped repeat regions.

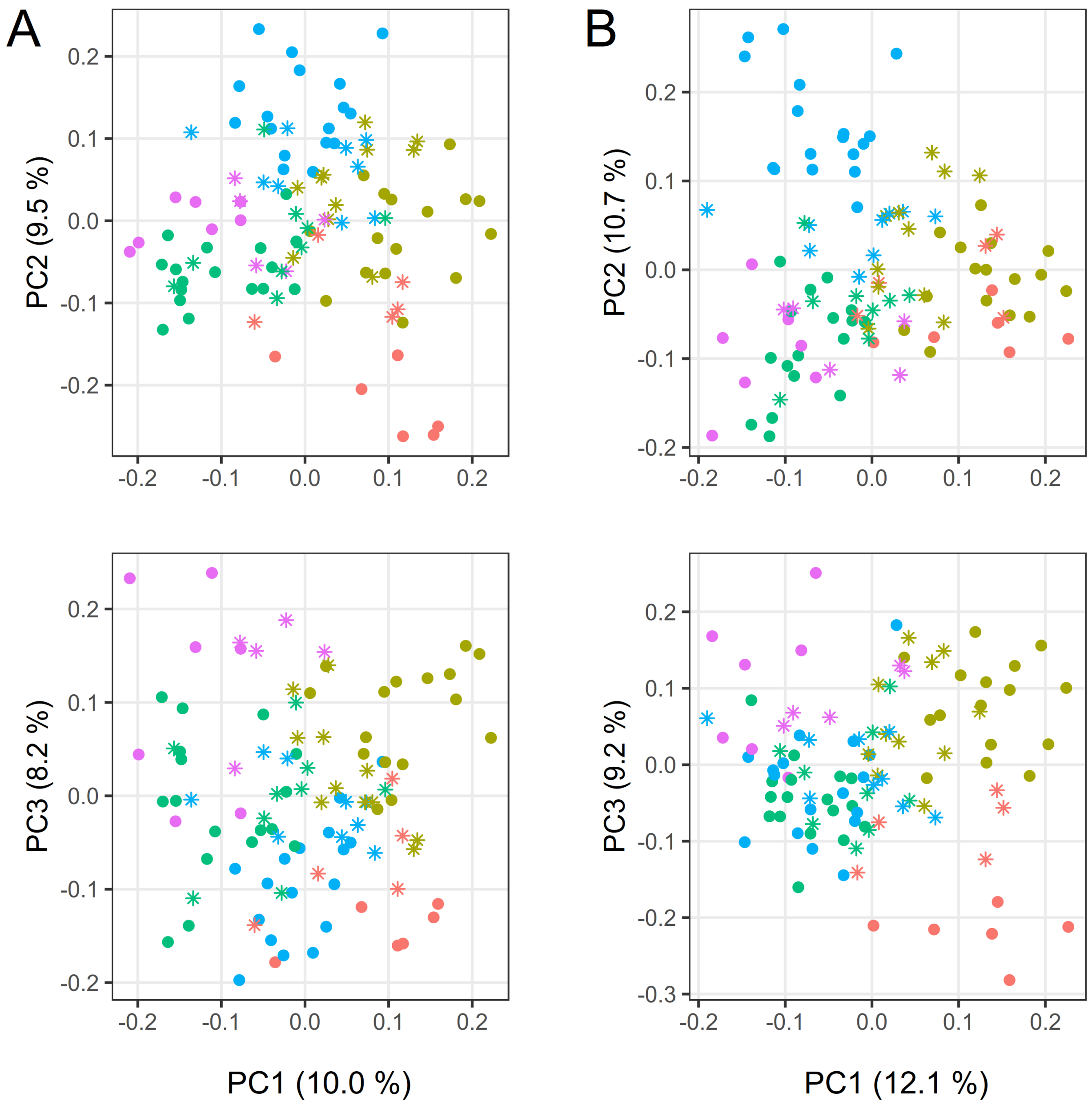

**Figure S17:** Population structure analyses of the whole Canadian soybean panel. PCA were computed based on (A) SVs discovered by Illumina and Oxford Nanopore and genotyped with Illumina data using Paragraph and (B) SNVs called by Platypus from Illumina data. The first three principal components are shown for each PCA. Samples are colored according to their population assignment by fastStructure based on the population with the highest q-value for each sample. A subset of the Illumina SNVs were used by fastStructure to infer the populations. Samples with a maximum q-value below 0.6 are considered admixed and are labeled with asterisks.

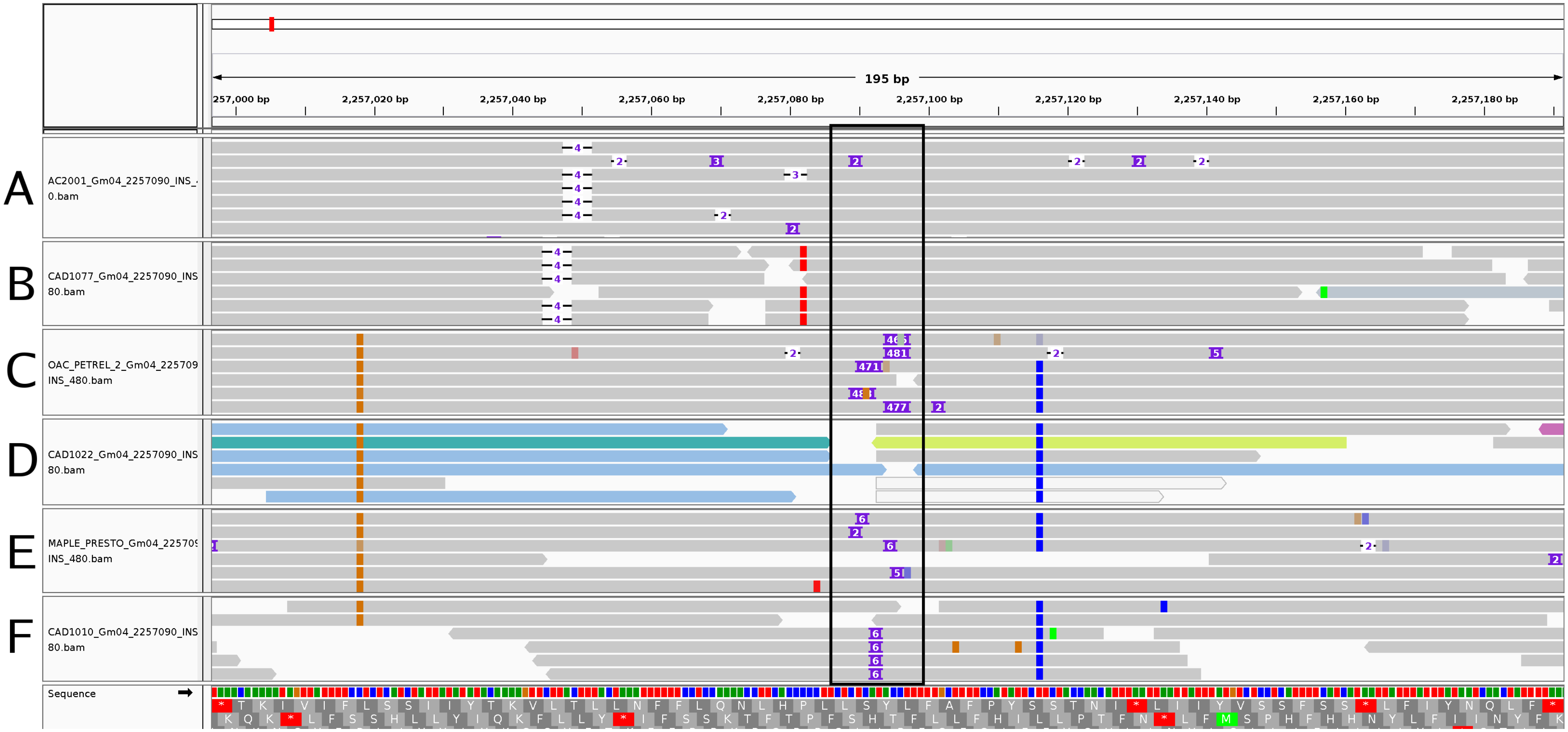

**Figure S18:** IGV screenshot of Oxford Nanopore and Illumina read alignments of three samples at the location of a polymorphic Stowaway TE insertion at Gm04:2,257,090. (A) Oxford Nanopore and (B) Illumina alignments of sample CAD1077/AC2001, which matches the reference (absence of TE insertion). (C) Oxford Nanopore and (D) Illumina alignments of sample CAD1022/OAC Petrel, which bears the 480-bp Stowaway TE insertion. (E) Oxford Nanopore and (F) Illumina alignments of sample CAD1010/Maple Presto, which bears a 6-bp insertion putatively due to excision of the TE. The black rectangle in the middle of the figure encloses the region where the insertion polymorphisms of interest occur.

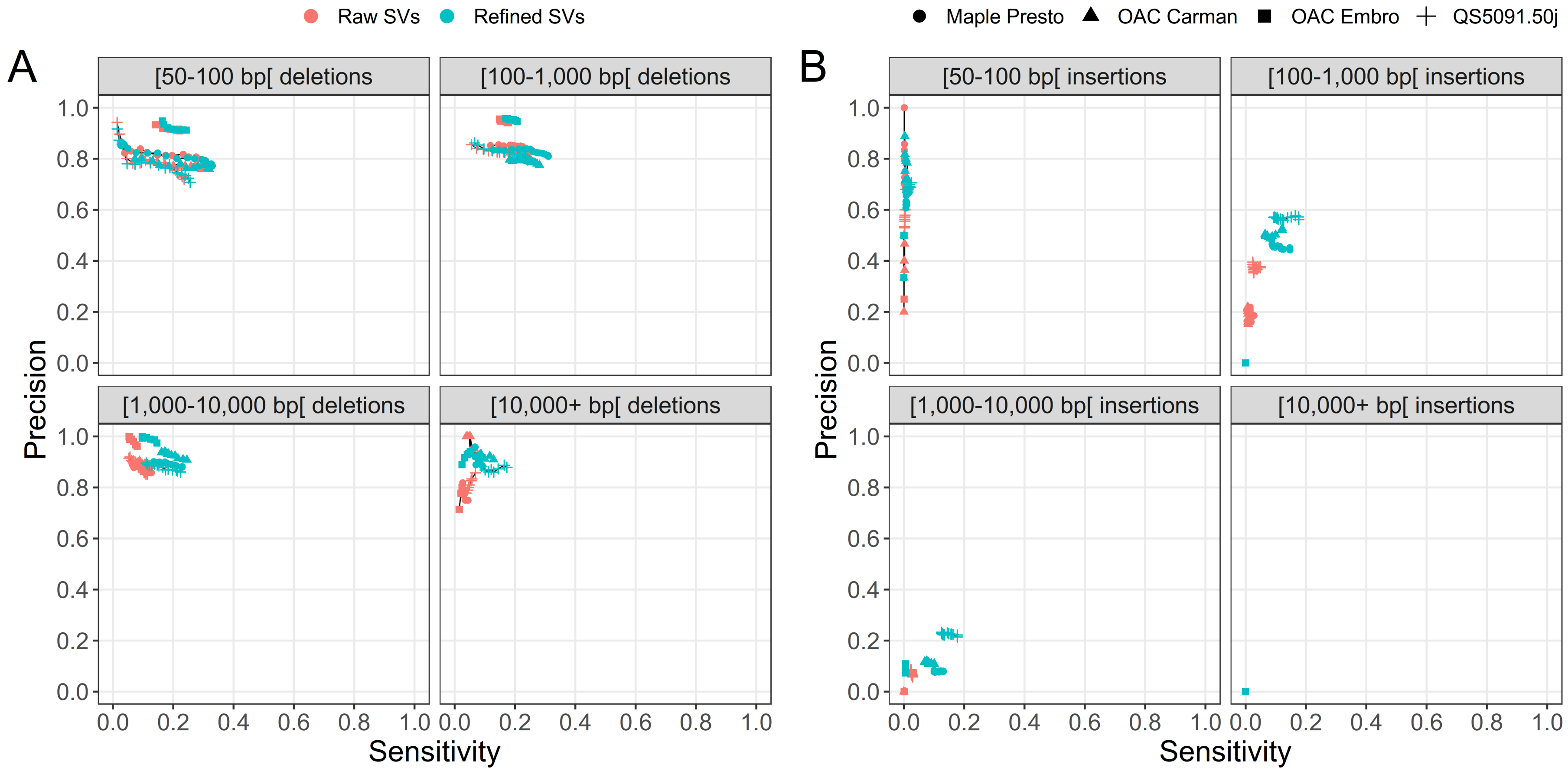

**Figure S19:** Genotyping sensitivity and precision of raw and refined (A) deletions and (B) insertions discovered by Sniffles on the Oxford Nanopore data for four samples when using BayesTyper as a genotyper. Each line represents data for a single sample whose SV breakpoints have either been refined (refined SVs) or not (raw SVs). The points correspond to different filtering thresholds on the genotype quality (GQ) required to support a genotype call.

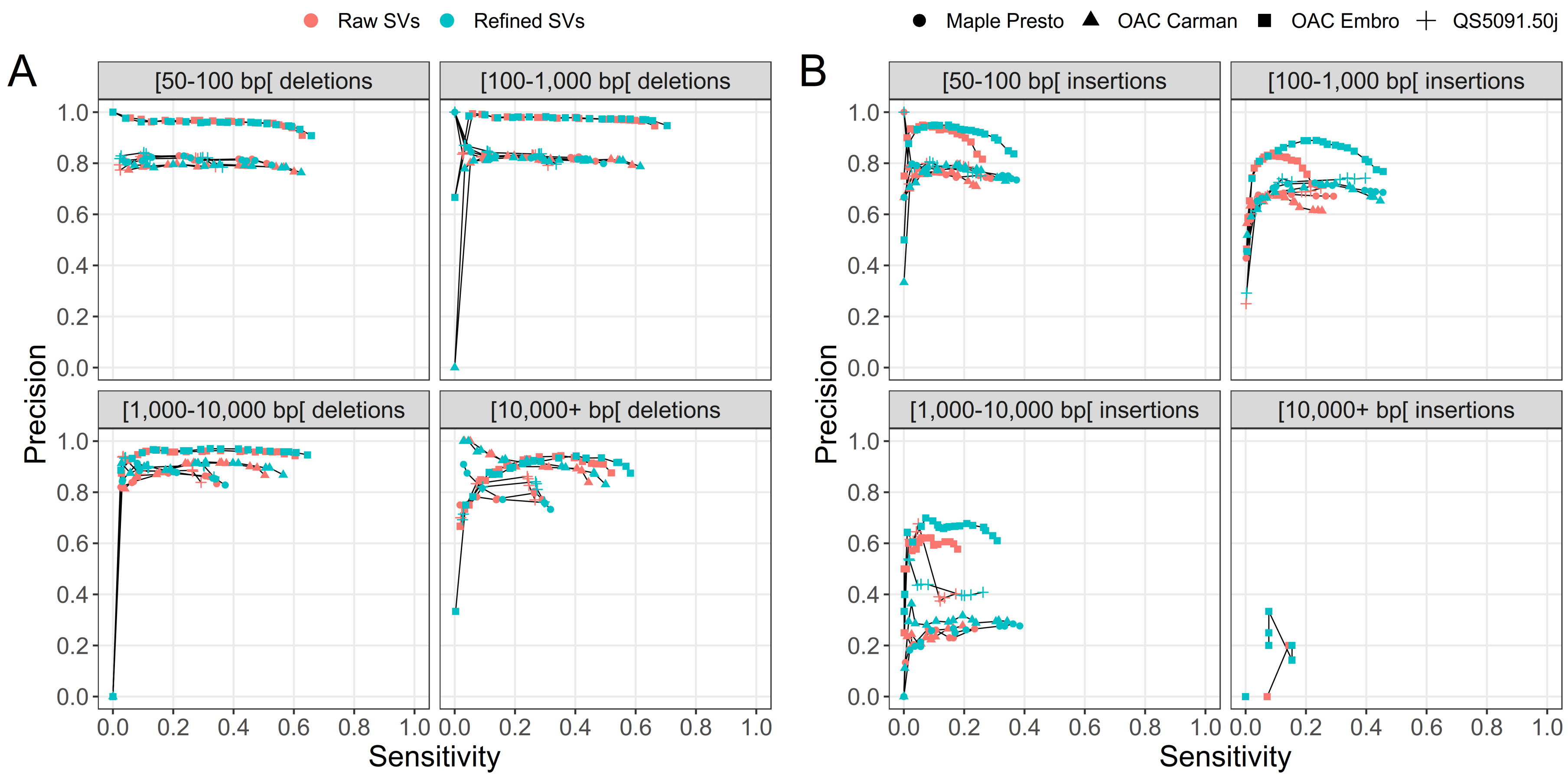

**Figure S20:** Genotyping sensitivity and precision of raw and refined (A) deletions and (B) insertions discovered by Sniffles on the Oxford Nanopore data for four samples when using vg as a genotyper. Each line represents data for a single sample whose SV breakpoints have either been refined (refined SVs) or not (raw SVs). The points correspond to different filtering thresholds on the genotype quality (GQ) required to support a genotype call.

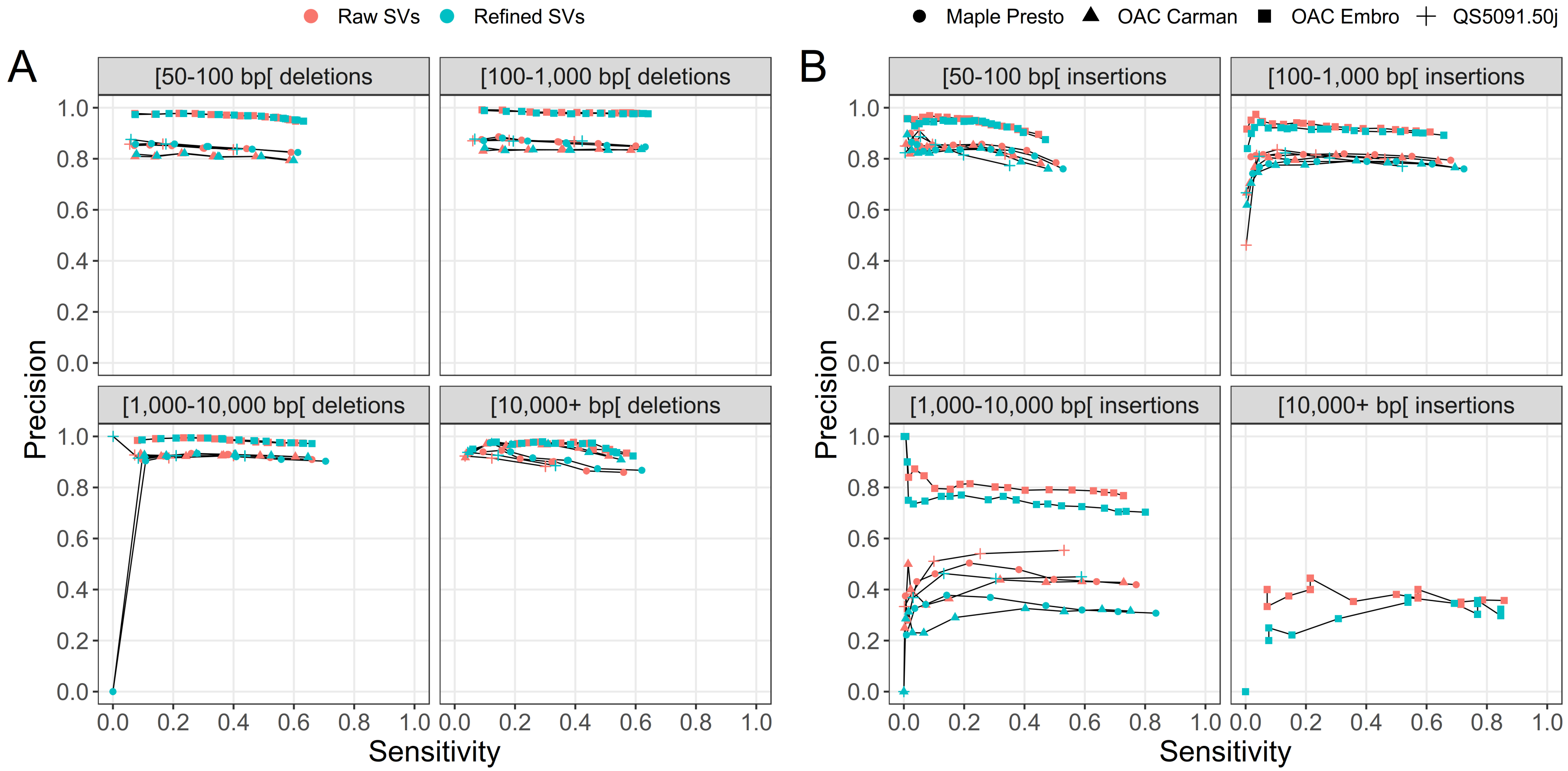

**Figure S21:** Genotyping sensitivity and precision of raw and refined (A) deletions and (B) insertions discovered by Sniffles on the Oxford Nanopore data for four samples when using Paragraph as a genotyper. Each line represents data for a single sample whose SV breakpoints have either been refined (refined SVs) or not (raw SVs). The points correspond to different filtering thresholds on the minimum number of Illumina reads required to support a genotype call.
